# Prolactin and DNA damage trigger an anti-breast cancer cell immune response

**DOI:** 10.1101/2020.08.31.274357

**Authors:** Ödül Karayazi Atici, Nayantara Govindrajan, Isbel Lopetegui Gonzalez, Constance A. M. Finney, Carrie S. Shemanko

## Abstract

There are conflicting reports on the role of prolactin (PRL) in breast cancer, and its role within the context of the tumour microenvironment is not well understood. In our previous study, we demonstrated a cross-talk between the ataxia telangiectasia-mutated (ATM) DNA damage response pathway and the PRL-Janus-kinase-2 (JAK2)-signal transducer and activator of transcription-5 (STAT5)-heat shock protein-90 (HSP90) pathway. To investigate the role of PRL in tumour initiation and the effect of DNA damage in vivo, we used a model of breast cancer initiation that assesses the ability of breast cancer cells to initiate orthotopic xenograft tumour formation after DNA damage. Breast cancer cells engineered to secrete human PRL or the control cells, were treated with the DNA damaging agent doxorubicin or vehicle and injected into the mammary fat pad of immune-deficient SCID mice. PRL secretion from human breast cancer cells did not change the tumour latency compared to controls, although combined doxorubicin and PRL treatment increased tumour latency. Depletion of glycolipid asialo ganglioside-GM1 positive immune cells using anti-asialo GM1 antibody resulted in faster tumour formation only in the PRL-secreting breast cancer cells that were pre-treated with doxorubicin, and not in the PRL-only or empty vector controls. Additionally, doxorubicin plus PRL treatment of breast cancer cells were shown *in vitro* to attract cytotoxic NK cells compared to controls, and that this was dependent on the PRLR. These results may shed light on the conflicting reports of PRL in breast cancer and demonstrate that combined breast cancer cell DNA damage and PRL exposure results in anti-tumour activity of asialo-GM1-positive immune cells.

## Introduction

Prolactin (PRL) is a peptide hormone that promotes proliferation, differentiation, survival and motility of mammary epithelial and mammary or breast tumour cells by inducing several signalling pathways upon binding to the PRL receptor (PRLR) (1). Although confirmed as a lactogenic hormone (2), its interaction with the PRLR is also implicated in the progression and metastasis of breast cancer (3–5). PRL is secreted from the pituitary gland, as well as from extrapituitary sites in humans where it behaves as a paracrine/autocrine signalling molecule (6, 7). Previous studies demonstrated that autocrine PRL in mammary specific neu-related lipocalin (NRL)-PRL transgenic mice contribute to increased mammary tumour formation (8–10).

PRL is a potential survival factor against apoptosis (11) and its proliferative effect is due to JAK2/MAPK activation (12). PRL-activated TEC/VAV/RAC pathway (13) and that of serine/threonine NIMA (never in mitosis A)- related kinase 3 (NEK3) is implicated in increased motility and invasion of breast cancer cells (14). Increased levels of autocrine PRL secretion from breast cancer cells were shown to induce proliferation of the cells, enhance PRLR expression and accelerate tumour growth (15). PRL has been demonstrated to have a role in cytotoxic resistance of breast cancer to a variety of chemotherapy drugs *in vitro* (16, 17).

In our previous study (Karayazi Atici et al. 2018), we demonstrated that PRL increases the viability of breast cancer cells treated with DNA-damaging agents, doxorubicin, or etoposide, and identified part of the molecular mechanism. The PRL-induced cellular resistance to DNA-damaging agents was specific to the PRLR and the DNA-damage response protein, ATM, was required for the PRL- Janus-kinase-2 (JAK2)-signal transducer and activator of transcription-5 (STAT5)- heat shock protein-90 (HSP90) pathway to mediate this resistance.

Transgenic mouse models indicate that PRL will promote tumour growth (8, 15, 18), however, there are some observations that PRL may have the opposite effect. MDA-MB-231 triple-negative breast cancer cells that were engineered to express the long form of the PRLR resulted in decreased tumour growth with recombinant PRL treatment in the non-obese diabetic/ severe combined immunodeficient (NOD)/SCID xenograft model (19). However, recent studies demonstrated isoform complexes formed by long PRLR mediate tumorigenic actions of PRL. While homomeric complexes of long PRLR promoted mammary differentiation, the intermediate and long PRLR heteromeric complexes promoted tumourigenesis (20). A novel PRL-humanized mouse model, NSG-Pro, emphasized the tumour-promoting role of endocrine PRL in PRLR-positive patient-derived xenograft models (21). These studies emphasize the complexity of PRL signalling and the need to understand and delineate the conditions by which PRL has differing effects on breast cancer.

In this study, we used an orthotopic model to investigate the role of autocrine PRL in breast tumour progression in the context of DNA damage and the presence of immune cells. SCID mice have severe deficiency in T and B cells, however, they have active natural killer (NK) cells, which sometimes limits the growth of human xenografts (Dorshkind 1985, Dewan et al., 2005). In order to deplete the activity of NK cells and asialo-GM1 positive immune cells in mouse models, an anti-asialo-GM1 antibody was developed (Kasai et al., 1981). Our experiments indicate that autocrine PRL supports the initiation and growth of xenografts, but that the combination of PRL and the DNA damage response triggers an attack from asialo-GM1-positive immune cells in SCID mice.

## Materials and Methods

### Materials

Doxorubicin (Sigma-Aldrich Canada Co., Oakville, ON, Canada), 10 mg, was dissolved in sterile DMSO in a final concentration of 100 mM and stored at - 20°C protected from light. Human recombinant PRL was purchased from A.F. Parlow, National Hormone and Pituitary Program, CA.

### Antibodies

For western blots following antibodies were used, monoclonal anti-human PRL antibody (R&D System, Minneapolis, MN, USA), rabbit anti-STAT5 (phospho Y694) primary antibody [E208] (Abcam), mouse STAT5 primary antibody (Transduction Laboratories, BD BioSciences, San Jose, CA, USA), rabbit anti-H3 primary antibody (Millipore, Billerica, MA, USA), anti-GRB2 (Transduction Laboratories, BD Biosciences, Clone 81).

### Cell culture and cell lines

Eight cell lines were used in this study. MCF7 (estrogen receptor+, p53 wild type), SKBR3 (HER2-positive, p53 mutant) human breast cancer cells, and NK92MI human NK cells were obtained and authenticated from American Type Culture Collection and were used within 6 months when revived from frozen storage.

Breast cancer cell lines were maintained in DMEM (Invitrogen, Burlington, ON, Canada), supplemented with 10% fetal bovine serum (FBS) (PAA Laboratories Inc., Etobicoke, ON), 100 µg/ml streptomycin and 100 units/ml penicillin, 2 mM L-glutamine (Invitrogen, Burlington, ON, Canada). Zeocin, 800 μg/ml (Invivogen, San Diego, CA, USA), was used for transfected MCF7 cells and all MCF7 cells were supplemented with 10 μg/ml insulin (BD Biosciences, Mississauga, ON). NK9MI cells were maintained in Minimum Essential Medium Eagle (MilliporeSigma, Burlington, ON, Canada) supplemented with 12.5% FBS, 12.5 Horse Serum (Invitrogen) 100 µg/ml streptomycin and 100 units/ml penicillin, 2 mM L-glutamine, 1 mM Sodium pyruvate (Invitrogen), 0.2 mM myo-inositol (MilliporeSigma), 0.02 mM folic acid (MilliporeSigma) and 0.1 mM B-mercapto- ethanol (Invitrogen). Cells were passaged up to a maximum of 30 passages post-expansion. Mycoplasma testing was routinely performed using pan-species primers and PCR.

### Creation of syngeneic lines

To prepare stable cell lines of MCF7 cells engineered to secrete autocrine PRL or carry the empty vector (EV), MCF7hPRL and MCF7 control (EV) cell lines, polyethyleminine (PEI) transfection was used. PEI MW25K (Polyscience Inc., Warrington, Pa, USA) was dissolved in water to a final stock concentration of 1 mg/ml. The human PRL coding sequence was cloned into pcDNA3.1/Zeo(+) mammalian expression vector (plasmid a gift from Dr. Vincent Goffin, Inserm and University Paris Descartes, Paris, France) (Liby et al. 2003); the expression of hPRL was driven by the cytomegalovirus promoter. Empty pcDNA3.1/Zeo(+) plasmid was prepared as a control for the human PRL plasmid. The stably transfected cells carrying the Zeocin antibiotic resistance gene (Sh ble) were selected with and maintained in Zeocin (800 μg/ml) (Invivogen, San Diego, CA, USA). Following 10-15 days of incubation in the presence of Zeocin, the colonies were selected with cloning cylinders (Fisher Scientific, Toronto, ON, Canada) using Dow Corning high-vacuum grease (Fisher Scientific, Toronto, ON, Canada). Cells carrying the empty vector were confirmed by PCR (Supplementary Method).

PRLR knockout (KO) SKBR3 cells were created using Clustered regularly interspaced palindromic repeats (CRISPR)/ CRISPR-associated protein 9 (Cas9) genomic engineering and the PRLR KO was confirmed by genome sequencing (Centre for Genome Engineering, University of Calgary). Wild-type (WT) and CRISPR Control (CC) (single-cell clone controls) cells were used as controls for two independent PRLR KO SKBR3 colonies (SKBR3KO1, SKBR3KO2).

### Cell viability assay

Alamar blue cell viability reagent (Invitrogen, Burlington, ON, Canada) was used to determine the cellular proliferation and viability of the cells. Cells were seeded into 96-well plates (5000 cells/well). The next day cells were treated with 1uM doxorubicin for 2 hours. The media was refreshed, and cell viability was measured after 48 hours of recovery time. Alamar blue reagent, 1/10^th^ of the total volume, was added directly to the cells in the culture medium. After 2.5 hours of incubation time, the fluorescence intensity was read using a plate reader with an excitation at 530 nm and an emission at 590 nm (Spectramax M4 Microplate Reader). The fluorescence intensity of doxorubicin-treated cells was divided by DMSO (vehicle) treated cells for normalization.

### Nuclear lysate extract

The nuclear lysate extract was used to extract phosphorylated-STAT5 (p- STAT5), STAT5 and Histone-H3 proteins, as previously described by (***22***). To determine p-STAT5 levels, 10^6^ cells were plated in 10cm plates, and the next day treated for 30 minutes with human recombinant PRL. Cells were scraped and lysed for nuclear protein extraction.

### Immunoblotting for STAT5, p-STAT5, Histone-H3, TBP-1, GRB2 and PRL

Immunoblotting for STAT5, p-STAT5 and GRB2, were as previously described (23). To detect Histone-H3, the PVDF membrane was wet with methanol and blocked in 3% dried skim milk in PBS overnight at 4°C, and washed three times with TBST 0.05%. The membrane was incubated with rabbit anti-H3 primary antibody (Millipore, Billerica, MA, USA) (1:1000 dilution) in 3% dried skim milk in PBS overnight at 4°C, washed and incubated in HRP conjugated goat anti-rabbit secondary antibody (1:10000 dilution) in 3% dried skim milk in PBS for 1 hour before chemiluminescence detection as described (24). To detect PRL, the membrane was blocked in 3% dried skim milk in TBST 0.1% for 2 hours, washed and incubated in monoclonal anti-human PRL antibody (R&D System, Minneapolis, MN, USA) (1:100 dilution) in 5% dried skim milk in TSBT 0.1% overnight at 4°C. The next day the blot was washed and incubated in HRP conjugated goat anti-mouse secondary antibody (1:10000 dilution) in 5% dried skim milk in TBST 0.1% for 30 minutes before washing and chemiluminescence detection.

### Secreted protein measurement from conditioned media

To investigate secreted PRL levels from MCF7hPRL, MCF7EV cells, 1x 106 cells were plated onto 10 cm plates. Conditioned media was removed from the cells at indicated time points or after 7 days, transferred into microcentrifuge tubes and centrifuged at 4°C at 13,200 rpm for 15 minutes. Centrifuged conditioned media containing secreting proteins was transferred into a new cold microcentrifuge tube, snap-frozen and stored at -80 °C until use (25). Conditioned media collection was started the day after doxorubicin treatment for treated cells and considered as day 1. The media was not refreshed for 7 days. Protein levels were determined by western blot or human PRL ELISA (Invitrogen, Burlington, ON, Canada). The manufacturer’s instructions were followed for the ELISA assay.

### Soluble Senescence-Associated b-galactosidase activity assay (ONPG assay)

Cells were plated onto 10 cm cell culture plates in 1x 106 cell number and the following day treated or not with human recombinant PRL (25 ng/ml) for 24 hours. The next day, cells were treated with vehicle control or doxorubicin (1 uM) for 2 hours and recovered 6 days with or without PRL. To prevent confluency- induced senescence, cells were split once if required during this 6-day recovery period. Cells were replated in 1 x 10^6^ cell numbers on the 4^th^ day of recovery. Media was aspirated at the end of 6 days and cells were washed twice with 1X PBS and trypsinized for 4 minutes at 37°C. Cells were collected within media and centrifuged at 800 rpm at 22°C for 4 minutes, the pellet was washed in 1X PBS and centrifuged one more time at 800 rpm at 22°C for 4 minutes. After removing the supernatant, the pellets were resuspended in 25 ul of 0.2 M phosphate buffer at pH 6.0 (87.7 ml of 0.4M sodium phosphate monobasic 12.3 ml of 0.4 M sodium phosphate dibasic heptahydrate, 100 ml ddH2O) and subjected to three cycles of freeze and thaw cell lysis in liquid nitrogen and 37°C water bath. Following lysis cycles, cells were centrifuged at 12,000 rpm at 4°C for 5 minutes and supernatant containing protein was transferred into a new microcentrifuge tube. The protein concentration was measured with BioRad protein assay and equal protein values were used for all experimental groups. The protein samples were incubated in 275 ul assay buffer containing 2 mM MgCl2, 100 mM β-mercaptoethanol, 1.3 mg/ml ONPG (from previously prepared 4 mg/ml 2-Nitrophenyl β-D-galactopyranoside in 0.2 M phosphate buffer pH 6.0), 0.2 M phosphate buffer pH 6.0 for 4 to 9 hours at 37°C. The reaction was stopped with 500 ul of 1M Na2CO3. Three aliquots of each reaction (200 ul) were transferred into a 96-well culture plate and absorbance was measured at 420 nm (Spectramax M4 Microplate Reader). The optical densities at OD 420 were averaged for internal and experimental replicates and standard deviations were calculated.

### Calcein-AM Assay

Calcein-AM assay was designed following the study by Somanchi et al. 2017. Calcein-AM (Invitrogen, ON, Canada) work solution (1 mg/ml in DMSO) was prepared in serum-free DMEM at 2 ug/ml concentration. Breast cancer cells were plated into 10 cm cell culture plates in 1x 10^6^ cell number, and the following day treated or not with human recombinant PRL (25 ng/ml) for 24 hours. The next day, cells were treated or not with doxorubicin (1 uM) for 2 hours and recovered for 48 hours. Breast cancer cells were washed twice with 1X PBS and trypsinized for 4 minutes at 37°C. Cells were collected within media and centrifuged at 500 rcf at 22°C for 4 minutes. After removing the supernatant, the cells were counted and resuspended in Calcein-AM-containing media (10^6^ cells/ml) and incubated for 30 min at 37° C. After staining with Calcein-AM, cells were centrifuged and washed twice with 1X PBS and resuspended in growth media. NK92MI human NK cells were collected and suspended in growth media. Calcein-AM stained breast cancer cells were seeded into 96-well plates and NK cells were added to breast cancer cells with 1:1 or 1:10 Target: Effector ratio.

Breast cancer cells and NK cells were co-cultured for 24 hours. After 24 hours, the media was gently mixed using a 200 ul tip and 150 ul from each well was transferred into a black-wall 96 well plate and the fluorescence was read at wavelength excitation: 485 nm, emission: 535 nm. Calcein-AM stained breast cancer cells without NK cells were used as a control for spontaneous release, and the breast cancer cells treated for 5 min with 1% Triton X-100 were used as maximum release control. The % lysis was calculated based on the calculation below. % lysis= [(Test release-spontaneous Release)/ (max release-spontaneous Release)] x 100

### Xenograft Animal Models

All animal procedures were carried out strictly following the Canadian Council for Animal Care guidelines and ethics approval from the University of Calgary Life and Environmental Sciences Animal Care Committee.

Nine-week-old Fox Chase SCID female mice (strain 236) were purchased from Charles River Laboratories (Montreal, QC, Canada). To deliver estrogen to the mice, 17β-Estradiol pellets (0.72 mg/pellet, 60-day release, Cat. No. SE-121) were (Innovative Research of America, Sarasota, Florida, USA) inserted subcutaneously. Three days after insertion of the pellet, breast cancer cells were injected into the 4th mammary fat pad as described below. The contralateral mammary fat pads were injected with PBS and Trevigen Cultrex BME mixture (Cedarlane, Burlington, ON, Canada).

MCF7 control and MCF7hPRL cells were plated and the following day indicated groups were treated or not with doxorubicin (1 μM) or vehicle for 2 hours and recovered for 48 hours. Cells were washed twice with 1 X PBS and trypsinized. Following centrifugation at 400 rpm for 4 minutes cells were resuspended in media and counted to obtain 1x 10^6^ cells per mouse. The counted cells were centrifuged at 400 rpm for 4 minutes at 4°C one more time and the pellets were washed with 1X PBS followed by the last centrifugation step at 400 rpm for 4 minutes at 4°C. The washed pellets were resuspended in 100 μl cold PBS and 100 μl Cultrex BME mixture. Cells were injected into the 4th mammary fat pad of mice based on the experimental groups. To deplete NK cell activity in SCID mice, 20 ul of the anti-mouse asialo-GM1 (Cedarlane, Burlington, ON, Canada) or control serum (ImmunoReagents, Raleigh, NC, USA) was injected intraperitoneally every 3-4 days for the duration of the experiment, following the titration data from the manufacturer. Tumour volumes were calculated as follows: [V = (W^2^ × L)/2], where V is tumor volume, W is tumor width, and L is tumor length (26).

### Mammary Gland Digestion

Mammary glands were resected and transferred to a sterile petri dish and minced using a sterile razor blade until a homogeneous tissue mixture was obtained with minimal cellular damage. The minced tissue was incubated with regular pipetting in a dissociation buffer containing gentle Collagenase/ Hyaluronidase (StemCell Technologies, Vancouver, BC, Canada) and DMEM/F12 (Gibco) supplemented with 10% FBS. The mixture was centrifuged at 400 rcf for 5 min at 4°C and the supernatant was discarded. The erythrocytes were lysed and removed using a 0.8% ammonium chloride solution in 1X PBS with 5% FBS. The solution was centrifuged at 400 rcf for 5 min at 4°C and the pellet was resuspended in 0.25% trypsin. The trypsin was deactivated by adding 1X PBS with 5% FBS and centrifuged. The pellet was resuspended in Dispase (5 U/ml, StemCell Technologies) and DNase I solution (1 mg/ml, StemCell Technologies). After pipetting, it was resuspended in 5% FBS containing 1X PBS and filtered using a 40 um strainer. The cells were directly used for flow cytometry analysis or fixed using 2% PFA after being stained with fixable viability stain.

### Flow Cytometry

Flow cytometry experiments were optimized with multiple experiments and the final experiments were performed with independent biological replicates. Each mammary gland or each experimental plate was considered as a biological replicate (n).

For the experimental glands that were collected 10 days after cell injection, 3 injected and uninjected mice were used, injected and contralateral glands were processed separately, isolated mammary cells were pooled for n=2 for injected and n=1 for uninjected group and directly used for flow cytometry staining and analysis.

For the anti-asialo GM1 and control serum injection experiments, 6 mice from each group were used to isolate mammary cells, after processing separately, isolated cells were pooled and stained for n=3 for each group.

For *in vitro* studies, 3 independent plates of cells were pre-treated or not with 25 ng/ml human recombinant PRL for 24 hours followed by 2 hours of doxorubicin treatment (1 uM). After 48 hours of recovery, the cells were collected and washed with 1x PBS and used for flow cytometry staining or fixed in 2% PFA after being stained with fixable viability stain. The independent biological replicates were represented as n=3.

Cells, 1 x 10^6^, were washed once and resuspended in sodium azide and protein-free Dulbecco’s Phosphate Buffered Saline (1X DPBS) without FBS. 0.3ul of BD Horizon™ Fixable Viability Stain eFluor450-A Stock Solution was added per 1 ml cell suspension and vortexed immediately. After 15 min dark incubation at room temperature, the cells were washed twice with FACS washing buffer supplemented with FBS. Cells were resuspended in FACS wash buffer, stained directly with antibodies or fixed in 2% PFA.

The cells were incubated with Mouse Fc Block purified anti-mouse CD16/CD32 mAb (BD Horizon) at 1 ug/million cells in 100 ul FACS buffer (5% FBS in PBS) for 5 min at 4°C. Master mixes containing fluorescent labelled antibodies, Florescence Minus One (FMO) controls, IgG and secondary antibody controls were prepared in FACS buffer; 100 ul of appropriate antibody mixture was added to the samples and incubated for 30 min in the dark. Samples were washed with FACS buffer and transferred to FACS tubes for immediate analysis.

The following FACS antibodies were used according to the manufacturer’s instructions: BV421 Rat Anti-Mouse F4/80 (BD Horizon^TM^ #565411), CD335 (NKp46) Monoclonal Antibody (29A1.4) PerCP-eFluorTM 710 (eBioscience, 46- 3351-82), CD49b (Integrin alpha 2) Monoclonal Antibody (DX5) (eBioscience, 14- 5971-85), CD45, HLA-ABC Monoclonal Antibody W6/32 (eBioscience, 14-9983- 82), Mouse IgG2a kappa isotype control (eBioscience, 14-4724-82), secondary antibody staining for HLA antibody Donkey anti-mouse IgG (H+L)-FITC (Jackson Immunoresearch, 715-095-151), CD155 Monoclonal Antibody (2H7CD155) FITC (eBioscience, 11-1150-42), mouse IgG1 kappa isotype control FITC (eBioscience, 11-4714-82), CD112 (Nectin-2) Monoclonal Antibody (R2.447) APC (eBioscience, 17-1128-42), Mouse IgG1 kappa isotype control APC (eBioscience, 17-4714-81), MICA/B Monoclonal Antibody (6DE) PE (eBioscience, 12-5788-42), mouse IgG2a kappa isotype control PE (eBioscience, 12-4724,82).

The FlowJo software was used for analysis of all FCS files. The viable cell population was detected with FVD eFluor 450 dye. Gates were set according to FMOs for each antibody and sample type.

### Gating Strategy

Single cells were gated based on forward and side scatter patterns. In cases where cell viability was checked, cells negative for fixable viability dye FVD eFluor 450 were selected as live cells. *In vivo*, the immune population was chosen by gating for CD45-AF700 positive cells. Within this CD45-positive population, NK cells were identified based on DX5-FITC positive staining, mature NK cells were identified based on NKP46-PerCP-efluor710 positive staining (27, 28), and macrophages were identified based on positive F4/80-BV421-A staining (Wilson et al. 2022). The values were evaluated in injected and contralateral glands, and normalized to the uninjected gland and presented as relative percentages of CD45+DX5+ cells, NKp46+ cells within CD45+DX5+cells, or CD45+F4/80+cells. Gates for each dye were adjusted based on single color controls, FMO controls and isotype staining for relevant antibodies.

### Statistics

Statistical significance was tested with a paired or unpaired Student t test, alternatively, analysis of variance (ANOVA) was used with post-testing for multiple comparisons. Log-rank (Mantel-Cox) and Gehan-Breslow Wilcoxon statistical analyses were used to determine differences in tumour latency. One-way ANOVA followed by Mann-Whitney U Test was used to analyze tumour volumes. Results were considered significant when *P* value was lower than .05 (*P*< .05).

## Results

### Preparation and validation of a PRL-secreting MCF7 cell line (MCF7hPRL)

In order to create a xenograft model whereby the human breast cancer cells would have a consistent supply of human PRL, MCF7 cells were stably transfected with a human PRL expression (hPRL) plasmid or the empty vector (EV). Cellular PRL levels were evaluated from whole cell extracts (Figure S1A). PRL secretion from transfected colonies was evaluated from conditioned media (CM), using western blot (Figure S1B). Two colonies (colony 1 and colony 5) with relatively high levels of PRL were pooled (MCF7hPRL) to use for future experiments. The MCF7EV control line was confirmed by amplification of the vector antibiotic (zeocin) resistance gene, Sh ble, in stable lines (Figure S1C). These syngeneic lines were used in subsequent experiments as a model for autocrine secretion from breast cancer cells (MCF7hPRL) and its control line (MCF7EV).

The autocrine PRL secretion was further evaluated from MCF7hPRL, MCF7EV and parental MCF7 cells over seven days from conditioned media without refreshing media, using an ELISA assay. MCF7hPRL cells were determined to secrete 24 ng/ml of PRL daily and the levels increased up to 38 ng/ml after seven days in CM. This is most likely due to increased cell number and cumulative PRL detected from CM over time. PRL secretion was not detected from parental MCF7 cells and a negligible amount of PRL secretion from MCF7EV cells detected at days 6 and 7 (0.24 and 0.52 ng/ml, respectively) demonstrated high variability (Figure 1A).

**Figure 1:**
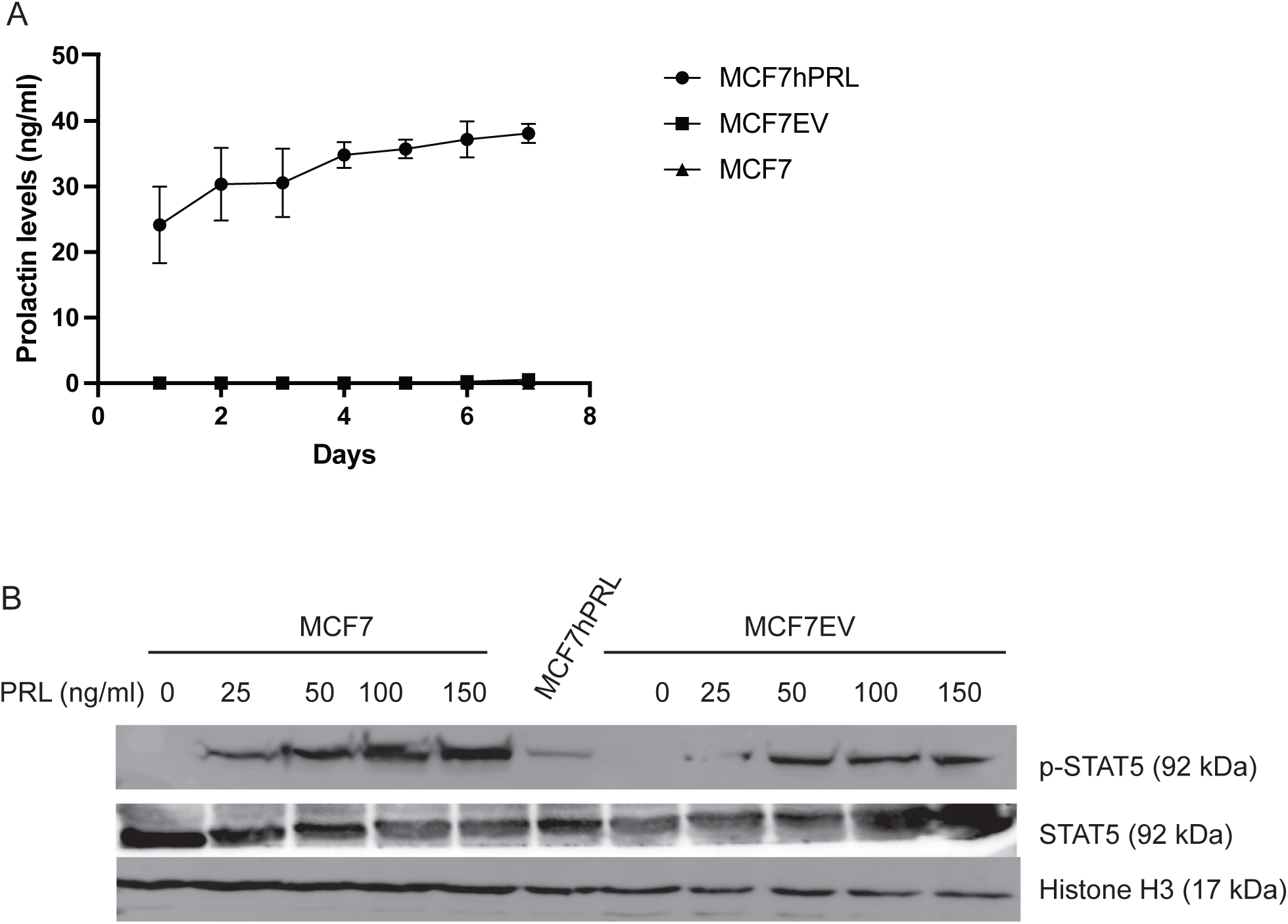
Confirmation of PRL-secreting and control lines. **A.** ELISA analysis of PRL secretion from MCF7hPRL, MCF7EV and parental MCF7 cells over 7 days. The conditioned media collection was started the day after plating the cells (1 x 10^6^ cells/10 cm plate). n=3 for each group. **B.** p-STAT5 and total STAT5 and histone-H3 (loading control) protein levels from untreated MCF7hPRL cells, or parental MCF7 and MCF7 control EV cells treated with increasing concentrations of human recombinant PRL.

In order to confirm that the autocrine PRL successfully activates the PRLR of the transfected MCF7 cells, we examined phosphorylated-STAT5 (p-STAT5), versus total STAT5 levels, as a read-out of PRL-JAK2-STAT5 pathway activation. Parental MCF7 and MCF7EV cells did not have detectable levels of p-STAT5 in the absence of PRL treatment, but both lines demonstrated p-STAT5 with increasing concentrations of recombinant human PRL (25 ng/ml, 50 ng/ml, 100 ng/ml and 150 ng/ml) (Figure 1B), confirming that the recombinant human PRL can activate STAT5. The MCF7hPRL cell line was shown to have levels of p- STAT5 most similar with the parental cell lines treated with 25 ng/ml PRL concentration, which is consistent with their calculated daily PRL secretion. Therefore, the MCF7hPRL cell line was confirmed to secrete PRL and the secreted PRL activates STAT5 as a readout of PRLR activation and the JAK2-STAT5 pathway.

### Autocrine PRL causes cellular resistance to DNA damage by doxorubicin

It has been observed previously that doxorubicin may impact PRL levels or PRL secretion (Howell et al., 2008). The effect of DNA damage by doxorubicin on PRL secretion was further evaluated by an ELISA assay from MCF7, MCF7EV and MCF7hPRL cells. PRL secretion was evaluated from CM over 7 days after 2 hours of doxorubicin treatment (1 uM) and increased PRL secretion was detected from MCF7hPRL cells, compared to untreated MCF7hPRL cells Figure 2A. The PRL secretion levels, however, decreased over 7 days of the period, which might be due to cell death induced by doxorubicin treatment. A negligible amount of PRL secretion was detected from MCF7EV or MCF7 cells treated or not with doxorubicin. Doxorubicin treatment does appear to initially increase PRL secretion from cells engineered to secrete hPRL but the effect may not be sustained.

**Figure 2.**
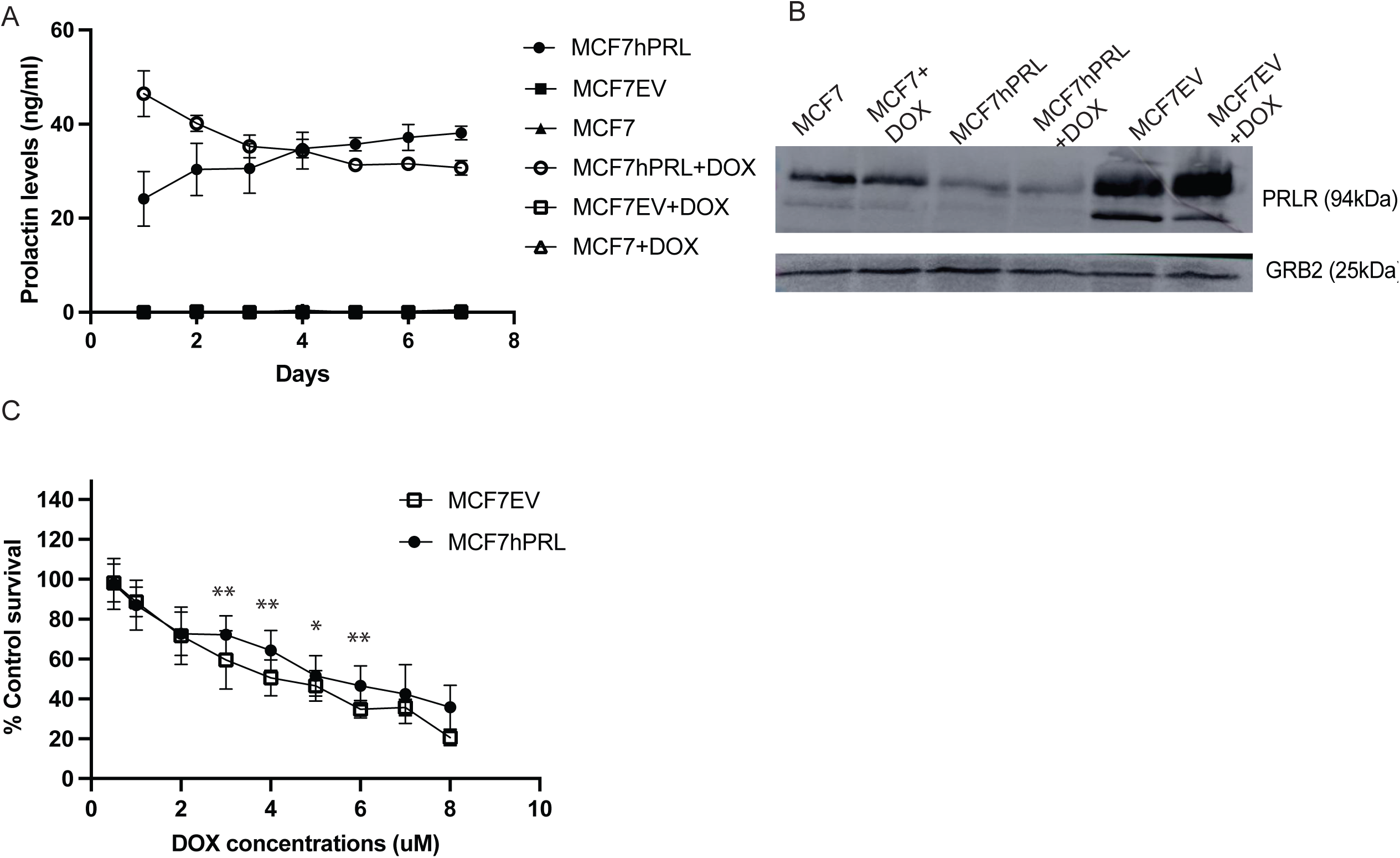
Autocrine PRL secretion causes resistance to DNA damage in vitro. **A.** ELISA analysis from MCF7hPRL, MCF7EV and parental MCF7 cells after doxorubicin treatment (1 uM-2 hrs treatment). The conditioned media collection was started the day after doxorubicin treatment. Secretion levels were presented as ng/ml, n=3 for each group. **B**. PRLR and GRB2 (loading control) protein levels from untreated and doxorubicin (1 uM- 2hrs) treated MCF7hPRL, MCF7EV and MCF7 cells. **C.** Cell viability (Alamar blue) assay showing autocrine PRL-mediated resistance to doxorubicin treatment in MCF7 Cells. Cells were treated with doxorubicin for 2 hrs, then recovered 48hrs. The viability of vehicle control (DMSO) treated cells was set to 100%. Graphs represent pooled experiments, n=12. Student t test was used for statistical analysis. Statistically significant analysis (*) denotes *P*<.05, (**) denotes *P*<.01. (Statistical differences for indicated DOX concentrations are, 3 uM (*P*=.006), 4 uM (*P*=.002), 5 uM (*P*=.04), 6 uM (*P*=.003).

To evaluate if doxorubicin treatment has any effect on PRLR levels, we examined the protein levels in the presence and absence of doxorubicin treatment. Our results demonstrated that MCF7EV cells had higher levels of PRLR, compared to parental MCF7 and MCF7hPRL cells. Interestingly MCF7hPRL cells showed lower levels of PRLR compared to the other two lines. Our results confirmed that doxorubicin did not affect PRLR levels in any of the MCF7 lines (Figure 2B).

In our previous publication (Karayazi Atici et al. 2018), we demonstrated that human recombinant PRL treatment increased the viability of breast cancer cells treated with doxorubicin. In this study, the cell viability was tested from MCF7EV and MCF7hPRL cells treated with increasing concentrations of doxorubicin. MCF7hPRL cells showed significant resistance to doxorubicin at 3, 4, 5 and 6 uM concentrations (Figure 2C), when compared to MCF7EV treatments. This result confirms that autocrine PRL also increases cellular viability against the DNA-damaging agent, doxorubicin.

### Autocrine PRL delays tumour latency in the presence of DNA damage

A novel orthotopic model of breast cancer was used to investigate the role of autocrine PRL and the DNA damage response on tumourigenicity and tumour volume. MCF7hPRL or control breast cancer cells were treated with the DNA damaging agent, doxorubicin (1 μM), and recovered for 48 hours before the cells were injected into the mammary fat pad of immune-deficient SCID mice. This examines the ability of cells to form a tumour after chemotherapeutic treatment of the cells, in a recurrence-style model. The dosage and length of treatment were based upon our previous work that demonstrated an active DNA damage response, specifically ATM phosphorylation (23). This model was used to test the effect of PRL on the tumour-initiating ability of breast cancer cells.

In order to test the effect of autocrine PRL on tumourigenicity, latency, and tumour size of breast cancer cells in the xenograft model, 500,000 MCF7hPRL or MCF7 control cells treated or not with doxorubicin, were injected into the number 4 mammary fat pad and monitored for 60 days. Tumour formation was observed as early as 10 days in mice injected with MCF7hPRL cells or doxorubicin-treated MCF7 cells. The majority of the mice carrying MCF7 control, doxorubicin-treated MCF7 cells and untreated MCF7hPRL formed palpable tumour by 20- or 30-days post-injection (Figure 3A). In contrast, only one mouse formed a palpable tumour on day 25 when injected with doxorubicin-treated MCF7hPRL cells, and no other detectable tumour formation was observed over 60 days in this group, which was significantly different than the MCF7hPRL injected group (Log-rank *P*=.039, Gehan- Breslow Wilcoxon *P*=.02). Therefore, there was a longer latency to tumour formation for PRL-secreting, doxorubicin-treated MCF7hPRL cells compared to all other conditions.

**Figure 3.**
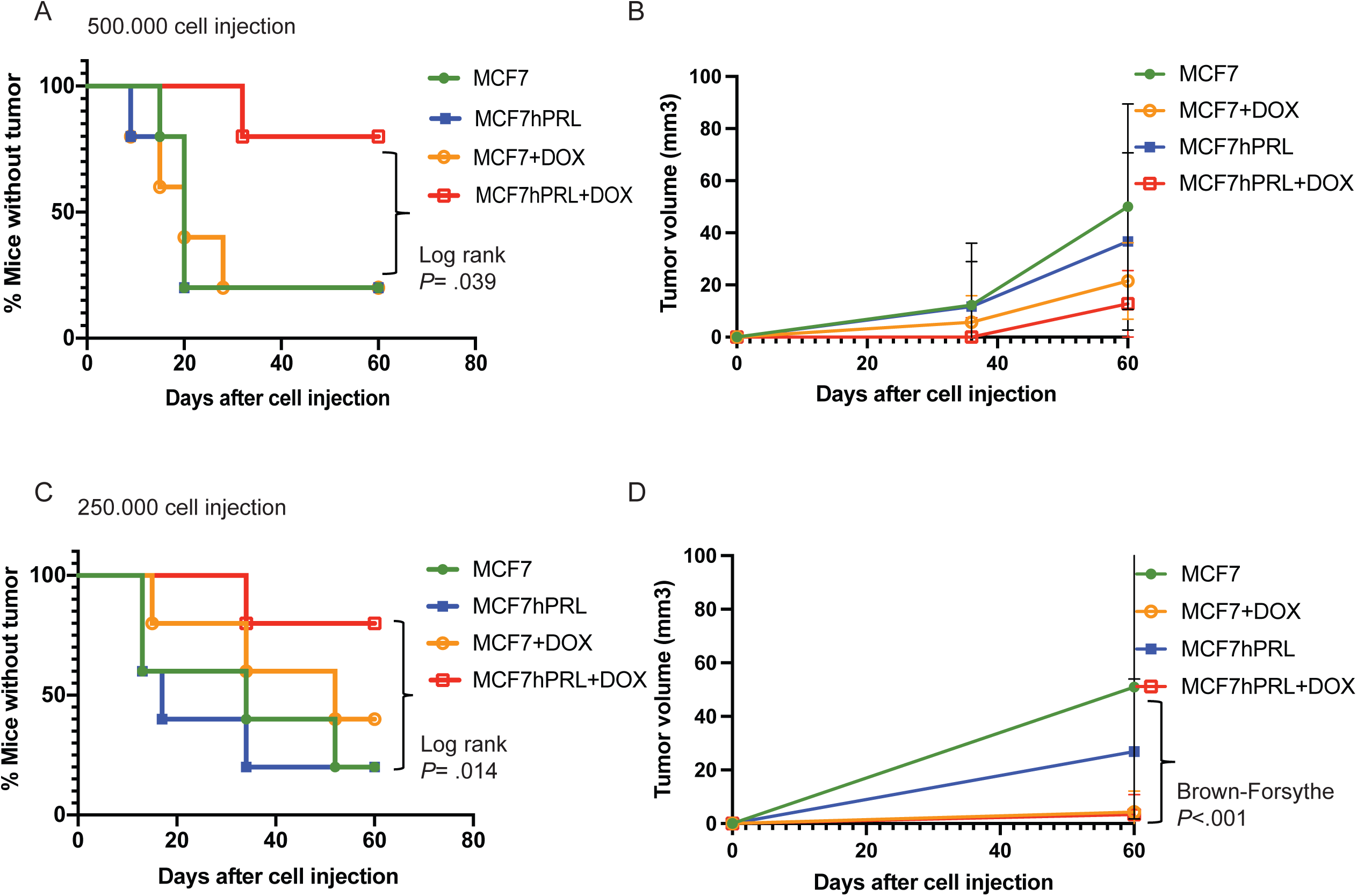
**Autocrine PRL delays tumour latency in the presence of DNA damage in SCID mice**. **A**. Tumour latency in SCID mice after injection of 500,000 MCF7 or MCF7hPRL cells -/+ doxorubicin over 60 days. **B**. Comparison of accumulated tumour volumes between treatment groups over 60 days. **C**. Tumour latency in SCID mice after injection of 250,000 MCF7 or MCF7hPRL cells -/+ doxorubicin over 60 days. **D**. Comparison of accumulated tumour volumes between treatment groups over 60 days. The sample size is n=5 mice for each group. Log-rank (Mantel-Cox) and Gehan-Breslow Wilcoxon tests were used for statistical analysis. Tumour volumes were tested using one-way ANOVA followed by Brown-Forsythe test.

According to the trend seen from the tumour volume graphs (Figure 3B), mice injected with MCF7 control and MCF7hPRL cells had the largest tumours, The mice injected with doxorubicin-treated MCF7 and doxorubicin-treated MCF7hPRL cells formed smaller tumours.

We repeated the above experiment but reduced the number of injected cells from 500,000 to 250,000 to be able to more subtly observe the latency time between the groups. The earliest palpable tumour formation was observed from day 13 and 15 after cell injection in the group of mice injected with MCF7 control, doxorubicin-treated MCF7 control and MCF7hPRL cells (Figure 3C), and majority of the mice formed tumour within 60 days post cell injection in these groups. Consistent with the first experiment, increased latency was also observed in the mice injected with doxorubicin-treated MCF7hPRL cells, as tumour formation started 34 days post cell injection, and only one mouse formed a tumour over 60 days (Figure 3C). There was a statistical difference between the group of mice injected with MCF7 and doxorubicin-treated MCF7hPRL cells (Log-rank *P*= .014, Gehan- Breslow Wilcoxon *P*= .027) and between the group of mice injected with MCF7hPRL and doxorubicin-treated MCF7PRL cells (Log-rank *P*= .042, Gehan- Breslow Wilcoxon *P*= .039) at 60 days (Figure 3C).

The tumour volume measurements demonstrated a trend that MCF7 cells formed the largest tumours, followed by MCF7hPRL cells, then doxorubicin- treated MCF7 cells. The tumours generated by doxorubicin-treated MCF7hPRL cells were very small in volume (Figure 3D), consistent with the previous experiment. According to the data analysis using Brown-Forsythe test, there was a statistical difference (*P*<.001) between tumours formed with MCF7 control cells and doxorubicin-treated MCF7hPRL cells (Figure 3D). When tumours were allowed to grow over 120 days, all animals acquired a tumour, and the delay in latency observed in doxorubicin-treated MCF7hPRL cells was not permanent (Figure S2A).

Overall, we observed that latency to tumour formation was significantly delayed in MCF7hPRL cells treated with doxorubicin in SCID mice and resulted in the smallest tumours at the end of the experiment, independent of cell number.

### Autocrine PRL and DNA Damage response attracts asialo-GM1 positive immune cells in SCID mice

In order to determine if the reason for delayed latency to tumour formation in the MCF7hPRL cells with doxorubicin-induced DNA damage was due in part to increased activity or the presence of immune cells, such as NK cells, we assessed the involvement of asialo-GM1 positive immune cells in the mechanism. We used the same recurrence animal model, with 500,000 cells in each group of MCF7 control or MCF7hPRL cells injected into SCID mice. Cells were treated, or not for 2 hours with doxorubicin (1 μM) followed by 48 hours recovery time in the absence of doxorubicin before injection, as before. Mice were injected with anti-asialo-GM1 antibody or control serum from the start of the experiment to address the effect on tumour initiation.

Importantly, anti-asialo-GM1 injection did not show any effect on the tumours in mice injected with doxorubicin-treated MCF7 control EV cells when compared with mice injected with control serum (Figure 4A). Tumour formation with MCF7hPRL cells occurred in 60 to 75% of animals after only 10 days and was also independent of anti-asialo-GM1 (Figure 4B). Tumour formation was, however, impacted when the MCF7hPRL cells were treated with doxorubicin. As expected, tumour formation of doxorubicin-treated MCF7hPRL cells was delayed in the animals injected with control serum. Tumour formation occurred on day 10 in 20% of mice injected with control serum, and 60% of the mice injected with anti-asialo GM1 antibody (Figure 4C). We confirmed our findings in a repeat experiment using 1 x 10^6^ MCF7hPRL cells treated with doxorubicin and 15 mice per group instead of 5 animals. While only 25% of mice formed a tumour by day 10 in the mice injected with doxorubicin-treated MCF7hPRL cells, anti-asialo GM1 injection resulted in tumour formation in 75% of the mice in 10 days (Log-rank *P*= .008, Gehan-Breslow Wilcoxon *P*= .008) (Figure 4D). Therefore, combined PRL and doxorubicin treatments were confirmed to result in delayed tumour initiation of breast cancer cells due largely to the action of asialo-GM1-positive immune cells. To rule out an effect of anti-asialo GM1 treatment on the viability of the breast cancer cells, we used an *in vitro* cell viability assay. MCF7, MCF7EV and MCF7hPRL cells were treated with estimated physiological levels of anti-asialo GM1 used in xenograft experiments, in the presence or absence of doxorubicin treatment. It was shown that anti-asialo GM1 did not have any effect on cell viability without or with doxorubicin, at 24 or 48 hours (Figure S3) or at 96 hours (Figure 4E, 4F). These results confirmed that the observed effects of anti-asialo treatment in mice were due specifically to the impact on anti-asialo GM1 positive cells.

**Figure 4.**
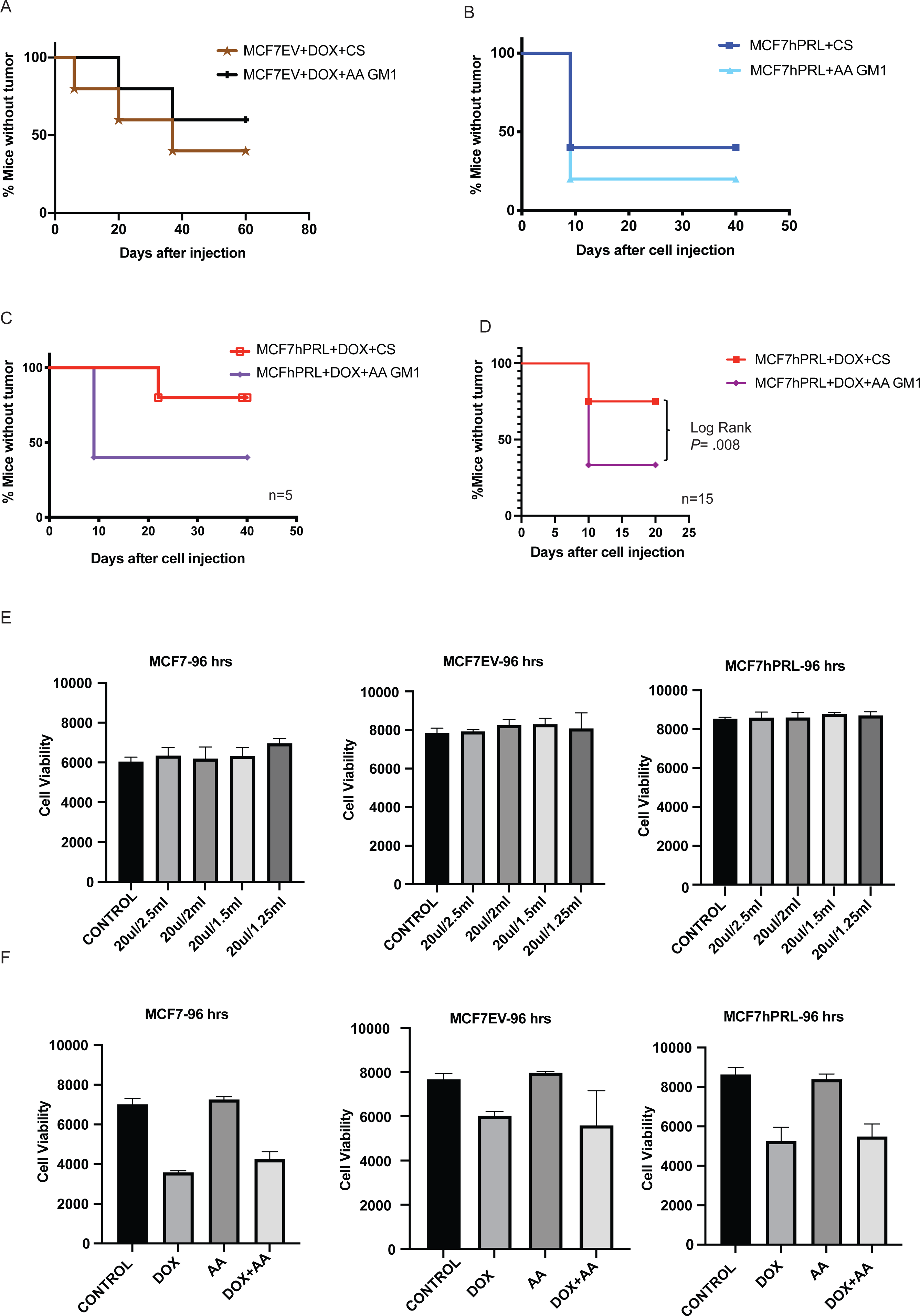
Autocrine PRL and DNA Damage response in breast cancer cells attract anti-asialo GM1 immune cells in SCID mice. **A.** Tumour latency after injection of 500.000 doxorubicin treated MCF7EV cells in mice treated with anti-asialo (AA) GM1 or control serum (CS). The sample size is n=5 mice for each group. **B.** Tumour latency after injection of 500,000 MCF7hPRL cells in mice treated with anti-asialo GM1 or control serum. The sample size is n=5 mice for each group. **C.** Tumour latency after injection of 500,000 doxorubicin treated MCF7hPRL cells in mice treated with anti-asialo GM1 or control serum. The sample size is n=5 mice for each group. Log-rank (Mantel-Cox) and Gehan-Breslow Wilcoxon tests were used for statistical analysis. **D.** Tumour latency after injection of 10^6^ doxorubicin treated MCF7hPRL cells in mice treated with anti-asialo GM1 or control serum. The sample size is n=15 mice for each group. Log-rank (Mantel-Cox) and Gehan-Breslow Wilcoxon tests were used for statistical analysis. **E**. Cell viability (Alamar blue) assay showing Anti-asialo GM1 does not affect the viability of MCF7, MCF7EV and MCF7hPRL cells. Cells were treated with 20 ul/2.5 ml, 20 ul/2 ml, 20 ul/1.5 ml, 20 ul/1.25 ml of Anti-asialo GM1. The cell viability was observed over 96 hrs. Graphs represent pooled experiments, n=6. **F**. Cell viability (Alamar blue) assay showing Anti- asialo GM1 does not affect the viability of MCF7, MCF7EV and MCF7hPRL cells in the presence and absence of doxorubicin. Cells were treated with 20 ul/2.5 ml, 20 ul/2 ml, 20 ul/1.5 ml, 20 ul/1.25 ml of Anti-asialo GM1. The cell viability was followed 96 hrs. Graphs represent pooled experiments, n=6.

### The NK and Macrophage recruitment changes after breast cancer cell and anti-asialo GM1 injections

It is known that anti-asialo GM1 antibody was originally designed to deplete NK cells (Yoshino et al. 2000), but it can also deplete other immune cells, such as macrophages (Wiltrout RH, 1985). It is also known that the DNA damage response attracts NK cells (29–31) We therefore examined the injected mammary glands to understand NK and macrophage recruitment, and depletion by the anti-asialo- GM1 antibody. To investigate the effect of DNA damaged breast cancer cells secreting autocrine PRL on immune cell trafficking, we injected 1 x 10^6^ doxorubicin treated-MCF7hPRL cells and after 10 days prepared the mammary glands for FACS analysis. We used CD45 as a nucleated hematopoietic marker for immune cells (Hermiston et al. 2003, Somasundara et al. 2021), DX5 (CD49b, Integrin alpha 2) as the marker for the general NK population in SCID mice, NKp46 as the major NK marker triggering antitumour activity and natural cytotoxicity (28, 32), and F4/80 as a macrophage cell marker (Cansever et al., 2023).

The recruitment of NK cells and macrophages was examined in both the injected and the contralateral glands after 10 days. The values were normalized to the uninjected gland and were presented as relative percentages of CD45+DX5+ cells, NKp46+ cells within CD45+DX5+ cells, or CD45+F4/80+ cells. The results showed an increasing trend in the recruitment of NK cells and macrophages, especially in the cytotoxic NKp46+ cells (Figure 5A, B, C).

**Figure 5.**
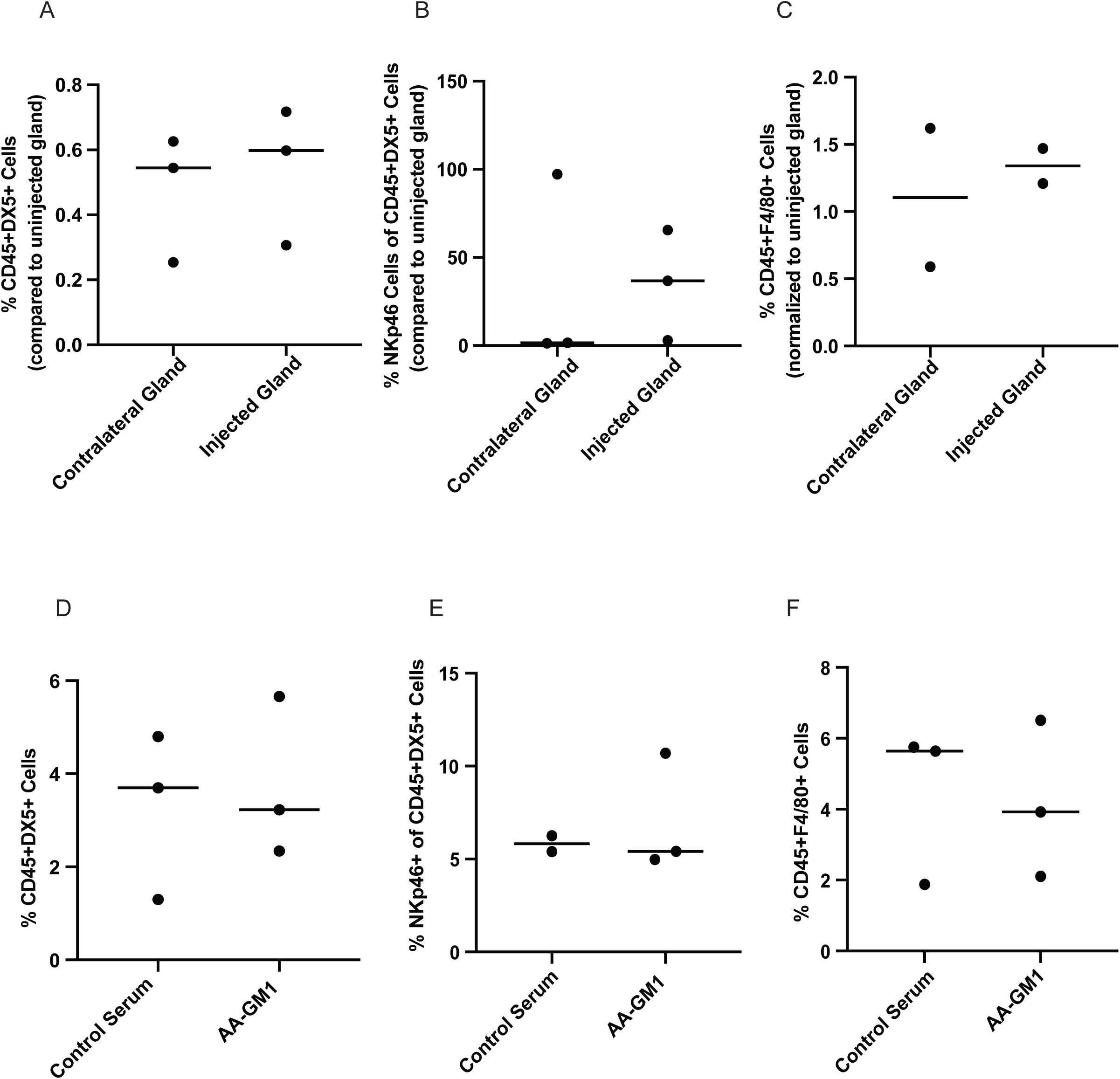
Percentage of NK cells and macrophages in SCID mice mammary gland after injecting DNA-damaged autocrine prolactin-secreting breast cancer cells. MCF7EV and MCF7hPRL cells were seeded and treated with 1uM doxorubicin for 2 hours the following day. After 48 hours of recovery time, 10^6^ cells were injected into SCID mice mammary glands. **A.B.C.** The mammary glands were collected after 10 days of cell injection and digested. and cells were analyzed using flow cytometry. Injected glands (n=3 mice), glands were pooled for staining (n=2), contralateral glands (n=3 mice), glands were pooled for staining (n=2) and uninjected mouse mammary gland (n=3 mice), glands were pooled for staining (n=1). **A.** Relative % of CD45+DX5+ cells. **B.** Relative % of Nkp46+ cells within CD45+DX5+ cells. **C**. %CD45+F4/80+ cells. For A-C, in all cases cell numbers were normalized to those seen in uninjected cells. **D.E.F.** The percentage of cells was evaluated from mammary glands after Anti-asialo GM1 and control serum treatment after 25 days. Anti-asialo GM1 group (n=6 mice), glands were pooled for staining (n=3), control serum group (n=6 mice), glands were pooled for staining (n=3). **D**. % CD45+DX5+cells. **E.** %NKp46+ cells within CD45+DX5+ cells. **F.** %CD45+F4/80+ cells.

The glands were also examined at the end of the anti-asialo GM1 injection experiment at day 25 using different mice instead of contralateral glands to distinguish the treatments. In this experiment, we compared the NK cells and macrophages between the mice that received control serum or anti-asialo GM1 treatment, after doxorubicin-treated MCF7hPRL cell injection. The results demonstrate that anti-asialo GM1 treatment caused a reduced trend in %CD45+DX5+ NK and %F4/80+ macrophage cell populations after 25 days of treatment (Figure 5D, E, F). We identified two NK populations and also macrophage cells in the mammary glands.

### DNA damage and PRL alter NK ligand levels in breast cancer cells

The major histocompatibility complex (MHC) known as human leukocyte antigen (HLA) class I molecules (HLA-A, -B, -C) are primary regulators of NK cell activation (Cruz-Tapias et al. 2013). Upon cellular stress and DNA damage, HLA class 1 levels may be downregulated, and activating receptors such as natural killer group 2D (NKG2D) and DNAX accessory molecule 1 (DNAM-1) are upregulated (Chan et al., 2014).

We evaluated HLA (-A, -B, and -C) levels from MCF7EV and MCF7hPRL by using FACS analysis. MCF7EV cells were pre-treated or not with human recombinant PRL (24hrs), followed by 2 hrs doxorubicin (1uM) treatment. MCFhPRL cells were treated with doxorubicin or not, and the HLA(-A,-B and -C) levels were measured by FACS analysis after 48 hours of recovery time. We determined a significant increase in median fluorescence intensity, particularly in doxorubicin-treated cells, in the presence of PRL. In MCF7EV cells, the HLA density was 1.3 fold higher in doxorubicin-treated cells compared to untreated (*P*=.0025) and 1.6-fold higher in doxorubicin+PRL-treated cells compared to PRL-treated control cells (*P*<.001). In MCF7hPRL cells, a 1.4-fold higher density was observed compared to untreated MCF7hPRL cells (*P*<.001) (Figure 6A). These results demonstrated that HLA class I molecules are upregulated by the DNA-damage response and also higher when also in the presence of PRL..

**Figure 6.**
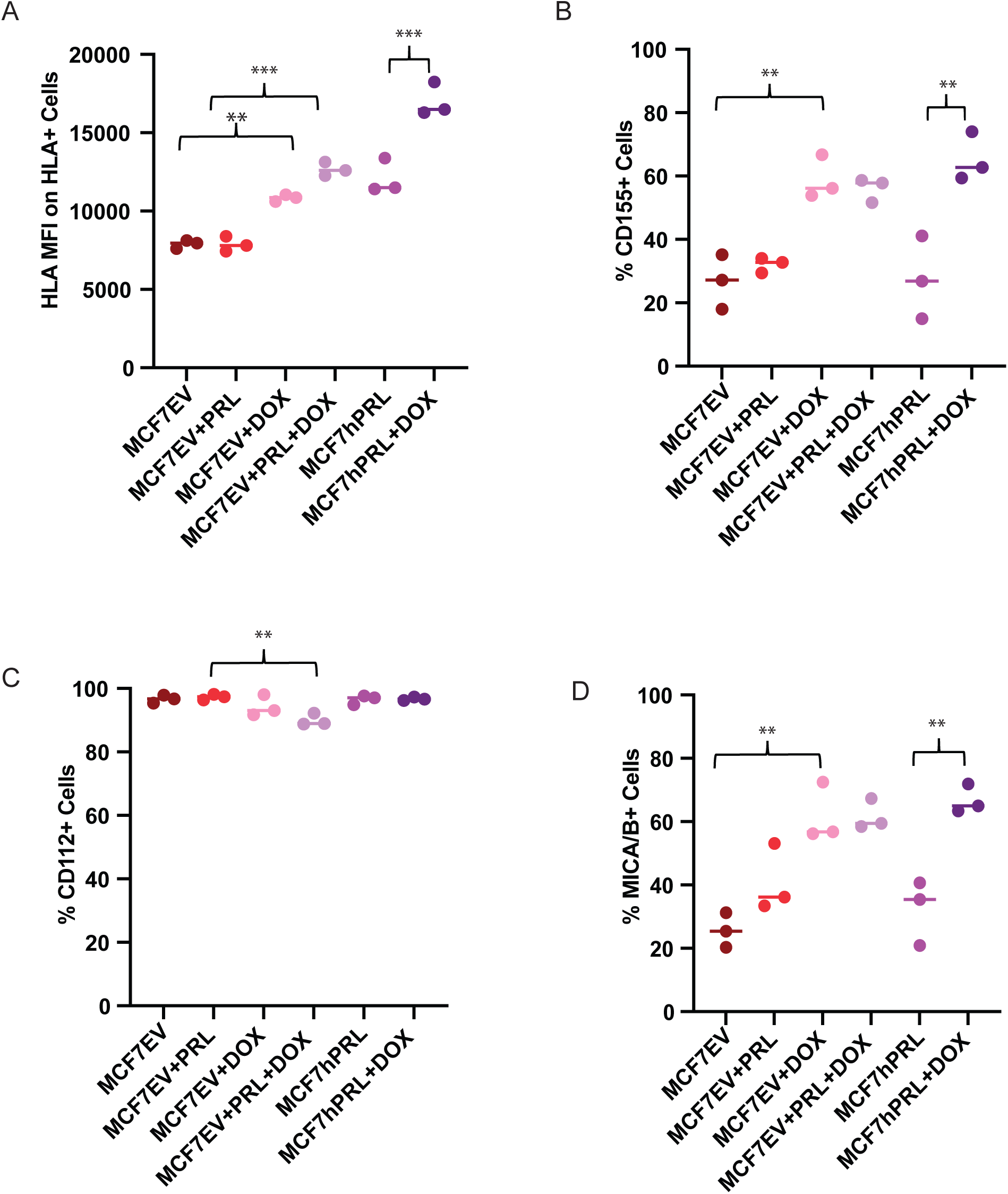
DNA damage and prolactin alters NK ligand expression in breast cancer cells. FACS analysis measuring the NK ligands from MCF7hPRL and MCF7EV cells in the presence or absence of doxorubicin treatment. Cells, 1x 10^6^, were seeded and MCF7EV cells were pre-treated with 25 ng/ml human recombinant prolactin for 24 hours, followed by 2 hours of doxorubicin treatment (1uM). Cells were trypsinized after 48 hours of recovery time and stained with CD155, CD112, MICA/B and HLA markers for FACS analysis. Graphs represent a pooled experiment, n=3. **A**. HLA median fluorescence intensity on HLA+ cells. **B**. %CD155+ cells **C**. % CD112+ cells **D**. %MICA/B+ cells. Statistically significant analysis (*) denotes *P*<.05, (**) denotes *P*<.01, (***) denotes *P*<.001.

To identify breast cancer ligands involved in triggering NK cell-activating receptors, we investigated protein levels of CD155, and CD112 as DNAM-1 ligands (Raulet et al., 2013, Bauer et al., 1999) and MICA/B as NKG2D ligand (Bottinno et al., 2003, Pende et al., 2005) from MCF7EV and MCF7hPRL cells by using FACS analysis under the same experimental conditions. In MCF7EV cells, the %CD155 levels showed a 2.1-fold increase with doxorubicin compared to the untreated control (*P*=.003) that did not change with the addition of PRL treatment (Figure 6B). In MCF7hPRL cells the increase was 2.4-fold higher with doxorubicin treatment compared to untreated cells (*P*<.001) (Figure 6B). Overall, the results demonstrated that NK ligand CD155 is upregulated in the presence of PRL and DNA damage, likely driven by doxorubicin treatment. The %CD112 levels demonstrated a decrease with PRL plus doxorubicin treatment compared to the PRL-treated (*P*=.003) MCF7EV cells (Figure 6C). The %MICA/B levels demonstrated a significant 2.4-fold increase in doxorubicin-treated MCF7EV cells (*P*=.001) compared to the untreated group, which was not increased by PRL (Figure 6C). Similar to CD155 levels, we observed an upregulation in MICA/B levels in doxorubicin-treated MCF7hPRL cells. There was a 2-fold increase in %MICA/B levels in doxorubicin-treated cells compared to untreated cells (*P*=.002) (Figure 6D). These results confirmed that some NK-activating ligands are upregulated in the presence of PRL and DNA-damage response, likely mostly driven by DNA damage.

### Autocrine PRL and DNA damage response increase senescence and NK cell- mediated killing

It is well known that DNA damage induces permanent cell cycle arrest, known as senescence. Cancer cells with wild-type p53, like MCF7 cells, undergo senescence in response to chemotherapy treatment (33). To investigate if senescence is part of the mechanism for increased latency and small tumour formation from doxorubicin-treated autocrine PRL-secreting cells, we measured senescence-associated beta-galactosidase levels.

As expected, doxorubicin treatment increased senescence significantly in MCF7 and MCF7EV cell lines compared to control cells (*P*=.008, *P*<.001). However, there was no significant difference detected with the addition of recombinant PRL in MCF7 and MCF7EV cells (Figure 7A, 7B). Doxorubicin treatment significantly increased senescence in MCF7hPRL cells compared to the control group (*P*<.001) (Figure 7C). This result confirmed that breast cancer cells go under DNA-damage-induced senescence likely independent of PRL.

**Figure 7.**
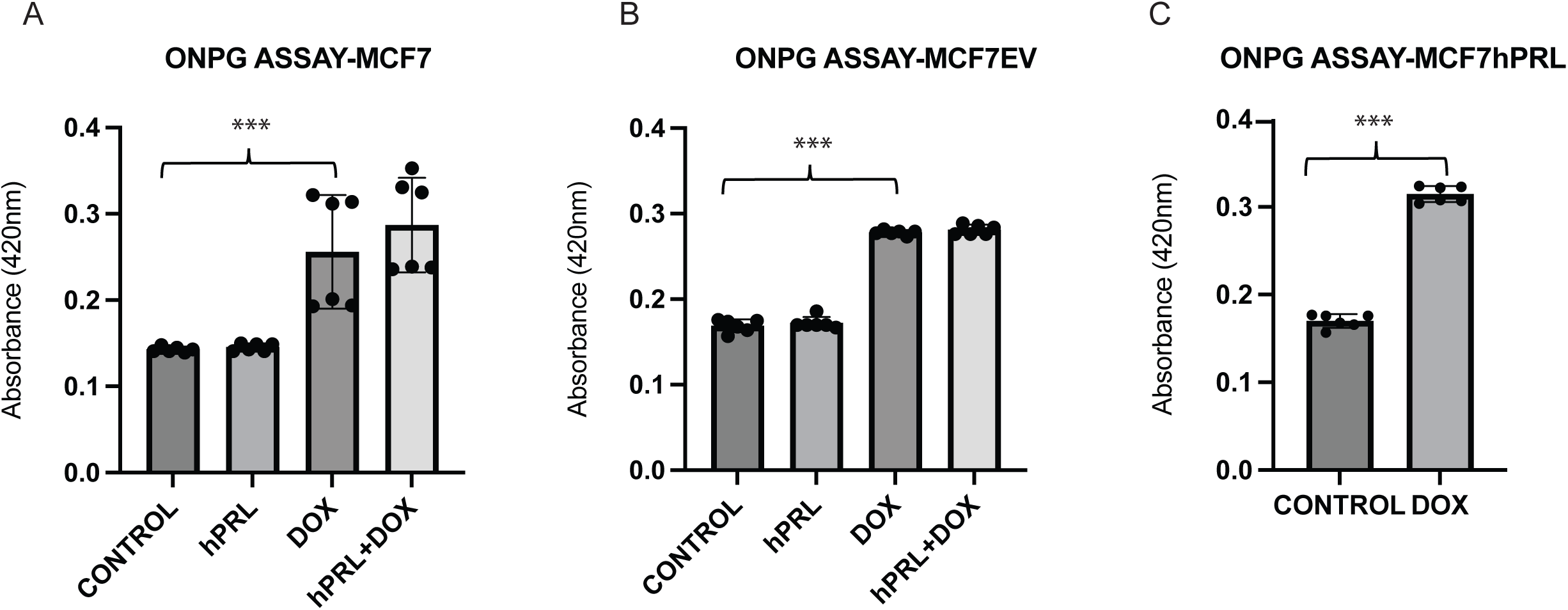
Autocrine prolactin and DNA damage increase cellular senescence A-B-C. Determining the increased effect of doxorubicin on senescence by ONPG assay in MCF7hPRL, MCF7EV and MCF7 cells. MCF7 and MCF7EV cells were pre-treated with human recombinant prolactin (25 ng/ml) for 24 hours, followed by 2 hours of doxorubicin treatment (1 µM). The cells recovered in the presence or absence of prolactin for 6 days. ONPG levels were measured by spectrophotometer. The graph represents 6 independent experiments. Statistically significant analysis (***) denotes *P*<.001.

Given that the anti-asialo GM1 antibody is well-known to deplete NK cells, and that we observed NK cells in the tumours, we investigated whether NK cell cytotoxicity may contribute to our observation of delayed latency after autocrine PRL and doxorubicin treatment. The attraction of NK cells by the DNA damage response and the presence of PRLR on NK cells were also considered during the experimental design. To examine if the combination of autocrine PRL secretion and DNA damage response makes cells more susceptible to NK cells, we performed a Calcein-AM assay to evaluate the percent lysis of breast cancer cells by NK cells under different experimental conditions. Human NK92MI NK cells were co-cultured with doxorubicin-treated and untreated MCF7, MCF7EV and MCF7hPRL at 10:1 and 1:1 effector: target ratios. To compare the effect of autocrine and recombinant PRL, MCF7 and MCF7EV cells were also treated with 25 ng/ml human recombinant PRL in indicated groups. The results demonstrated that DNA damage increases NK-mediated lysis significantly in all three MCF7 lines at 1:1 (Figure S4) and 10:1 ratios (Figure S4D, Figure 8 A, B). In MCF7EV cells, human recombinant PRL and doxorubicin treatment increased the percentage of lysis 5-fold compared to doxorubicin alone treatment (*P*= .001) (Figure 8A). The strongest effect was observed in autocrine PRL-secreting cells treated with doxorubicin, with the percentage of lysis 133-fold higher compared to untreated control (*P*<.001) and 220-fold higher compared to vehicle-treated control (*P*<.001) (Figure 8B). The results confirmed that doxorubicin and PRL, particularly autocrine PRL, increases the susceptibility of breast cancer cells to the cytotoxicity of NK cells.

**Figure 8.**
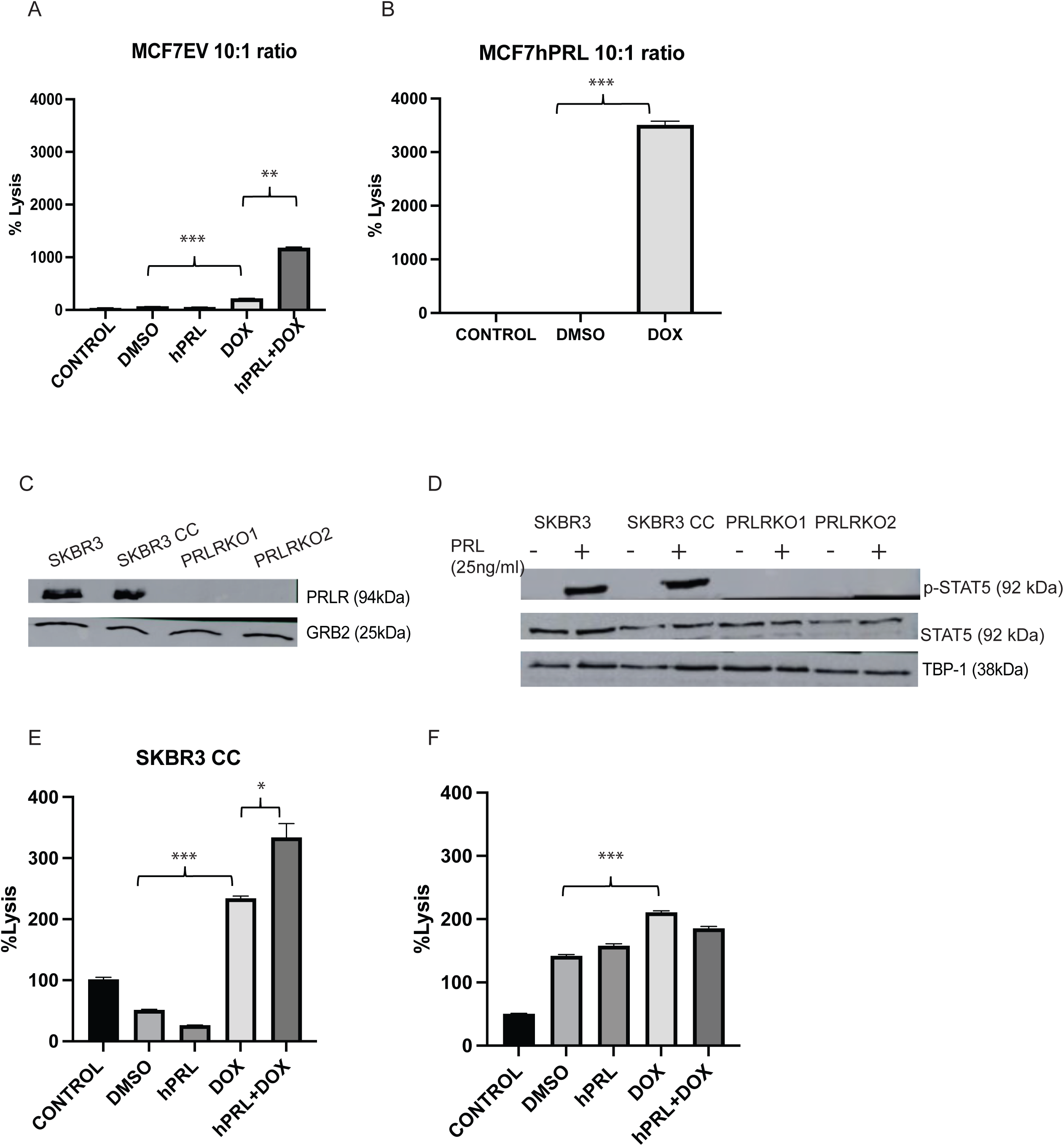
Autocrine prolactin and DNA damage increase NK cell-mediated cell lysis *in vitro*. A-B. Calcein-AM assay determining the NK cell-mediated lysis of MCF7hPRL and MC7EV cells in the presence or absence of DNA damage. MCF7EV cells were pre-treated with human recombinant prolactin (25 ng/ml) for 24 hours, followed by 2 hours of doxorubicin treatment (1 uM). Cells were trypsinized after 48 hours of recovery time and co-cultured with NK cells in a 10:1 effector/target ratio. The cell viability of breast cancer cells was determined by Calcein-AM assay and the %Lysis was calculated. C. PRLR and GRB2 (loading control) protein levels from SKBR3, SKBR3 CC, SKBR3PRLRKO1 and SKBR3PRLRKO2 cells. D. p-STAT5 and total STAT5 and TBP-1 (loading control) protein levels from SKBR3, SKBR3 CC, SKBR3PRLRKO1 and SKBR3PRLRKO2 cells treated or not with 25 ng/ml human recombinant PRL. E. Calcein-AM assay determining the NK cell-mediated lysis of SKBR3CC (Crispr Control cells) cells in the presence or absence of DNA damage at 1:10 effector/ target ratio. F. Calcein-AM assay determining the NK cell-mediated lysis of a PRLR knockout cell clone (PRLRKO) in the presence or absence of DNA damage at 1:10 effector/ target ratio. Statistically significant analysis (*) denotes *P*<.05, (**) denotes *P*<.01, (***) denotes *P*<.001.

### Active PRLR and DNA damage response increase NK cell-mediated killing

To further evaluate if the observed effect is due to the PRLR, we performed additional Calcein-AM assays with SKBR3 cells that have a CRISPR/Cas9 mediated deletion of the *PRLR* gene, SKBR3PRLR-knockout-(KO). The western blots confirm the presence of PRLR in the parental and CRISPR Control (CC) SKBR3 cells, whereas no PRLR was detected in two *PRLR* KO SKBR3 lines (SKBR3PRLRKO1 and SKBR3KO2) (Figure 8C). As expected, there was no STAT5 phosphorylation in the *PRLR* KO lines compared to SKBR3 and SKBR3- CC cells treated with 25 ng/ml human recombinant PRL (Figure 8D).

We performed Calcein-AM assays in co-cultures with NK92MI cells and SKBR3 cell lines at a 10:1 effector: target ratio (Figure S4F, Figure 8E, F). The results demonstrated similar results as with MCF7 cell lines, although the SKBR3 cell lines were more resistant to lysis overall compared to the MCF7 lines.

In SKBR3 CC cells, doxorubicin treatment increased the NK-medicated lysis significantly compared to vehicle control (*P*<.001), and prolactin plus doxorubicin increased lysis significantly compared to PRL (*P*=.005) or doxorubicin alone (*P*=.01) or (Figure 8E). Using the SKBR3PRLR KO line, although doxorubicin-induced lysis by NK cells, there was no PRL-mediated increase in lysis in the doxorubicin-treated cells, indicating the effect of PRL was PRLR-mediated. In the SKBR3PRLR KO cells, doxorubicin significantly increased NK lysis compared to vehicle control (*P*<.001) (Figure 8F). We confirmed increased NK-mediated lysis with doxorubicin treatment in SKBR3, SKBR3 CC, and SKBR3 PRLRKO cells. PRL plus doxorubicin treatment resulted in a further increase in NK-mediated lysis in SKBR3CC cells, and this was abrogated by loss of the PRLR.

## Discussion

In our breast cancer recurrence model, we have demonstrated that autocrine PRL in the tumour microenvironment does in fact not delay tumour initiation, but in the context of the DNA damage response, combined PRL signaling with DNA damage leads to immune cell attack and increases the latency of tumour formation. These observations may clarify conflicting observations on the role of PRL in breast cancer tumourigenesis and tumour progression.

The role of autocrine PRL in tumour biology has been previously studied in three different animal models. Production of PRL under the control of the metallothionein gene promoter in transgenic mice resulted in mammary tumour formation in 100% of the females (18). In a transgenic mice model that overexpresses PRL within mammary epithelial cells, under the control of the neu- related lipocalin NRL promoter (3), autocrine PRL was demonstrated to induce mammary tumours in NRL-PRL transgenic mice (8, 9). In a study from Liby and colleagues (15), MDA-MB-435 cells, whose origin may be melanoma or melanoma-like breast cancer cells (34), were genetically engineered to overexpress PRL and the cells were injected into the mammary fat pad of nude mice. PRL-secreting cells were shown to increase tumour growth 2-4-fold when compared with parental MDA-MB-435 cells. Additionally, tumours formed with PRL-secreting cells were demonstrated to metastasize to the lymph nodes. Autocrine PRL was overall shown to promote tumour initiation, tumour growth and metastasis in *in vivo* models.

PRLR function in tumour progression or metastasis has also been studied *in vivo* via delivery of recombinant PRL to animal models. Tumours generated in mice using 7,12-Dimethylbenz[*α*]anthracene (DMBA) regressed with the PRLR antagonist, LFA102, a humanized neutralizing monoclonal antibody directed against the extracellular domain of PRLR (35). Using a splice-modulating oligomer that knocked down only the long PRLR form *in vivo*, the long PRLR form was shown to be responsible for lung and liver metastasis and cell survival in primary mammary syngeneic tumours of 4T1 cells and BT-474 xenografts in *in vivo* models of breast cancer (Yonezawa et al., 2015). Overall, these models indicate that PRL and the PRLR support breast tumour progression and metastasis.

In our xenograft model, the autocrine PRL-secreting breast cancer cells or control cells were pre-treated with doxorubicin or vehicle and injected into the mammary fat pad of SCID mice, and the tumourigenicity and tumour volume were observed over time. The doxorubicin pre-treated PRL-secreting cells formed tumours later when compared with control, PRL-secreting, or doxorubicin-only treated cells. The tumour volumes were also of smaller size in the PRL and doxorubicin-treated groups. In the immune-deficient SCID mice used in this study, T and B cells are deficient, however NK cells, macrophages and basophils are present (Dorshkind 1985, Kumar et al., 1989, Dewan et al., 2005). In order to investigate the mechanism and the effect of the microenvironment on tumourigenicity we used the anti-asialo GM1 antibody, traditionally used to deplete NK cells. The anti-asialo GM1 antibody shortened the time to tumour formation of doxorubicin pre-treated PRL-secreting cells, which implicated that the combination of PRL secretion and DNA damage attracts immune cells that interfere with tumour initiation and/or progression.

In a conflicting report, PRL was observed to suppress tumourigenicity of triple-negative breast cancer cells in the non-obese diabetic NOD/SCID mouse xenograft model. The NOD-SCID model has a deficiency in T and B cells and reduced NK cell activity (36). MDA-MB-231 triple-negative breast cancer cells were stably engineered to express the long-form PRLR using a doxycycline-dependent expression system (19). In this inducible model, both doxycycline and PRL were provided simultaneously at the time of tumour formation. We speculate that the observed PRL-mediated reduction of tumour growth might be due to a small population of anti-asialo GM1 immune cells activated by the combination of PRL and accumulated DNA damage from doxycycline (37). Alternatively, the PRLR- transduced MDA-MB-231 cell line may be the biological difference resulting in their observations.

We previously demonstrated that ATM was required for PRL-JAK2-STAT5- induced clonogenic survival and cell viability after DNA damage from doxorubicin (23). We did not observe greater tumour size in this *in vivo* setting due to the action of asialo-positive immune cell components in the tumour microenvironment in the presence of both PRL and DNA damage. We hypothesize that the *in vivo* mechanism that triggers immune cell attack also involves the cross-talk of the PRL and DNA damage response pathways in breast cancer cells.

The anti-asialo antibody was designed to deplete NK cells, but previous studies have discovered that the anti-asialo antibody can also target basophil cells (Nishikado et al. 2011), NK, NKT, CD8+T, γδT, some CD4+T cells, macrophages, and eosinophils (38–42). Although SCID mice do not have functional B or T cells, they do have functional NK cells, macrophages, eosinophils and basophils (43), of which NK cells (44), eosinophils (45), basophils (46) and macrophages (Komohara et al., 2023) have been implicated in anti-tumour activity.

NK cells are cytotoxic lymphocytes of the innate immune system and have a critical role in immune response to tumours and viral infections. DNA damage has been demonstrated to activate several ligands specific for NK cell receptors (reviewed in (47). Chemotherapeutic agent-induced cellular stress and DNA damage response do attract NK cells (29–31) and increase ligand expression of NK cell receptors such as NK2GD and DNAM-1 in ATM and ATR-dependent manner (48). Doxorubicin-mediated DNA damage was shown to increase NK cells and T cell-mediated killing of tumour cells in a mechanism that involves TRAIL receptor signaling (49).

PRL has been implicated to have an important role in the recruitment of immune cells to the mammary gland. PRL treatment of adult female black 6 mice increased lymphocyte number in the mammary gland. Experiments with normal mammary HC11 cells *in vitro* revealed that PRL increased the migration of B cells, CD4+ T cells, CD4+ memory T cells, CD8+ memory cells, macrophages, monocytes, neutrophils, eosinophils (50) and basophils (Nishikado et al. 2011, Freeman et al., 2000). It was not reported if PRL recruited NK cells to mammary epithelial cells. PRL was also shown to directly improve the antitumour effects of NK cells in Balb/c and SCID mice via the PRLR on NK cells (51–53). PRL has been shown to directly upregulate the NK major receptors involved in lytic cell death (54). In our study, we found that breast cancer cells have increased levels of certain ligands that activate major NK receptors in the presence of PRL and DNA damage response. This could be part of the mechanism of the observed antitumor effect in SCID mice.

In summary, we observed that PRL supports mammary tumour formation, but that PRL in combination with the DNA damage response, increases tumour latency of breast cancer cells in an orthotopic xenograft model in a mechanism that involves asialo-GM1-expressing immune cells, likely NK cells. This may have implications for the use of PRLR antagonists during anti-cancer treatment with doxorubicin.

## Acknowledgements

CSS was responsible for study conception and design, OKA, NG, ILG for the acquisition of data, CSS, OKA, NG, ILG, CAMF for experimental design, analysis and interpretation of data, and all authors contributed to the writing of the manuscript. We would like to acknowledge the service provided by Centre for Genome Engineering and Flow Cytometry Facility at the University of Calgary.

## References

1. Shemanko CS. Prolactin receptor in breast cancer: marker for metastatic risk. J Mol Endocrinol. 2016;57(4):R153–R65.

2. Horseman ND, Zhao W, Montecino-Rodriguez E, Tanaka M, Nakashima K, Engie SJ, Smith F, Markoff E, Dorshkind K. Defective mammopoiesis, but normal hematopoiesis, in mice with a targeted disruption of the prolactin gene EMBO J. 1997;16(23):6926–35.

3. Arendt LM, Schuler LA. Transgenic models to study actions of prolactin in mammary neoplasia. Journal of mammary gland biology and neoplasia. 2008;13(1):29–40.

4. Sutherland A, Forsyth A, Cong Y, Grant L, Juan TH, Lee JK, et al. The Role of Prolactin in Bone Metastasis and Breast Cancer Cell-Mediated Osteoclast Differentiation. Journal of the National Cancer Institute. 2016;108(3):djv338.

5. Yonezawa T, Chen KH, Ghosh MK, Rivera L, Dill R, Ma L, et al. Anti-metastatic outcome of isoform-specific prolactin receptor targeting in breast cancer. Cancer letters. 2015;366(1):84–92.

6. Ben-Jonathan N, Mershon JL, Allen DL, Steinmetz RW. Extrapituitary prolactin: distribution, regulation, functions, and clinical aspects. Endocr Rev. 1996;17(6):639–69.

7. Muthuswamy SK. Autocrine prolactin: an emerging market for homegrown (prolactin) despite the imports. Genes & development. 2012;26(20):2253–8.

8. Arendt LM, Rugowski DE, Grafwallner-Huseth TA, Garcia-Barchino MJ, Rui H, Schuler LA. Prolactin-induced mouse mammary carcinomas model estrogen resistant luminal breast cancer. Breast Cancer Res. 2011;13(1):R11.

9. Rose-Hellekant TA, Arendt LM, Schroeder MD, Gilchrist K, Sandgren EP, Schuler LA. Prolactin induces ERalpha-positive and ERalpha-negative mammary cancer in transgenic mice. Oncogene. 2003;22(30):4664–74.

10. O’Leary KA, Shea MP, Schuler LA. Modeling prolactin actions in breast cancer in vivo: insights from the NRL-PRL mouse. Advances in experimental medicine and biology. 2015;846:201–20.

11. Perks CM, Keith AJ, Goodhew KL, Savage PB, Winters ZE, Holly JM. Prolactin acts as a potent survival factor for human breast cancer cell lines. Br J Cancer. 2004;91(2):305–11.

12. Yamauchi T, Yamauchi N, Ueki K, Sugiyama T, Waki H, Miki H, et al. Constitutive Tyrosine Phosphorylation of ErbB-2 via Jak2 by Autocrine Secretion of Prolactin in Human Breast Cancer. J Biol Chem. 2000;275(43):33937–44.

13. Miller SL, DeMaria JE, Freier DO, Riegel AM, Clevenger CV. Novel association of Vav2 and Nek3 modulates signaling through the human prolactin receptor. Mol Endocrinol. 2005;19(4):939–49.

14. Miller SL, Antico G, Raghunath PN, Tomaszewski JE, Clevenger CV. Nek3 kinase regulates prolactin-mediated cytoskeletal reorganization and motility of breast cancer cells. Oncogene. 2007;26(32):4668–78.

15. Liby K, Neltner B, Mohamet L, Menchen L, Ben-Jonathan N. Prolactin overexpression by MDA-MB-435 human breast cancer cells accelerates tumor growth. Breast cancer research and treatment. 2003;79(2):241–52.

16. Idelman G, Jacobson EM, Tuttle TR, Ben-Jonathan N. Lactogens and estrogens in breast cancer chemoresistance. Expert review of endocrinology & metabolism. 2011;6(3):411–22.

17. LaPensee EW, Ben-Jonathan N. Novel roles of prolactin and estrogens in breast cancer: resistance to chemotherapy. Endocrine-related cancer. 2010;17(2):R91–107.

18. Wennbo H, Gebre-Medhin M, Gritli-Linde A, Ohlsson C, Isaksson OG, Tornell J. Activation of the prolactin receptor but not the growth hormone receptor is important for induction of mammary tumors in transgenic mice. J Clin Invest. 1997;100(11):2744–51.

19. Lopez-Ozuna VM, Hachim, I.Y., Hachim, M.Y., Lebrun, J.J., Ali, S. . Prolactin Modulates TNBC Aggresive Phenotype Limiting Tumorigenesis Endocrine-related cancer. 2019;26(3):321–37.

20. Grible JM, Zot P, Olex AL, Hedrick SE, Harrell JC, Woock AE, et al. The human intermediate prolactin receptor is a mammary proto-oncogene. NPJ Breast Cancer. 2021;7(1):37.

21. Sun Y, Yang N, Utama FE, Udhane SS, Zhang J, Peck AR, et al. NSG- Pro mouse model for uncovering resistance mechanisms and unique vulnerabilities in human luminal breast cancers. Sci Adv. 2021;7(38):eabc8145.

22. Perotti C, Karayazi O, Moffat S, Shemanko CS. The bone morphogenetic protein receptor-1A pathway is required for lactogenic differentiation of mammary epithelial cells in vitro. In Vitro Cell Dev Biol Anim. 2012;48(6):377–84.

23. Karayazi Atici O, Urbanska A, Gopinathan SG, Boutillon F, Goffin V, Shemanko CS. ATM Is Required for the Prolactin-Induced HSP90-Mediated Increase in Cellular Viability and Clonogenic Growth After DNA Damage. Endocrinology. 2018;159(2):907–30.

24. Perotti C, Liu R, Parusel C, Böcher N, Schultz J, Bork P, et al. Heat shock protein 90alpha (Hsp90alpha), a prolactin-Jak2-Stat5 target gene identified in breast cancer cells, is involved in apoptosis regulation. Breast Cancer Research. 2008;10(6):R94.

25. Howell SJ, Anderson E, Hunter T, Farnie G, Clarke RB. Prolactin receptor antagonism reduces the clonogenic capacity of breast cancer cells and potentiates doxorubicin and paclitaxel cytotoxicity. Breast Cancer Res. 2008;10(4):R68.

26. Faustino-Rocha A, Oliveira PA, Pinho-Oliveira J, Teixeira-Guedes C, Soares-Maia R, da Costa RG, et al. Estimation of rat mammary tumor volume using caliper and ultrasonography measurements. Lab Anim (NY). 2013;42(6):217–24.

27. Krneta T, Gillgrass A, Chew M, Ashkar AA. The breast tumor microenvironment alters the phenotype and function of natural killer cells. Cell Mol Immunol. 2016;13(5):628–39.

28. Miao M, Masengere H, Yu G, Shan F. Reevaluation of NOD/SCID Mice as NK Cell-Deficient Models. Biomed Res Int. 2021;2021:8851986.

29. Raulet DH, Gasser S, Gowen BG, Deng W, Jung H. Regulation of ligands for the NKG2D activating receptor. Annual review of immunology. 2013;31:413–41.

30. Raulet DH, Guerra N. Oncogenic stress sensed by the immune system: role of natural killer cell receptors. Nature reviews Immunology. 2009;9(8):568–80.

31. Soriani A, Iannitto ML, Ricci B, Fionda C, Malgarini G, Morrone S, et al. Reactive oxygen species- and DNA damage response-dependent NK cell activating ligand upregulation occurs at transcriptional levels and requires the transcriptional factor E2F1. J Immunol. 2014;193(2):950–60.

32. Abel AM, Yang C, Thakar MS, Malarkannan S. Natural Killer Cells: Development, Maturation, and Clinical Utilization. Front Immunol. 2018;9:1869.

33. George N, Joshi MB, Satyamoorthy K. DNA damage-induced senescence is associated with metabolomic reprogramming in breast cancer cells. Biochimie. 2024;216:71–82.

34. Korch C, Hall EM, Dirks WG, Ewing M, Faries M, Varella-Garcia M, et al. Authentication of M14 melanoma cell line proves misidentification of MDA-MB- 435 breast cancer cell line. Int J Cancer. 2018;142(3):561–72.

35. Damiano JS, Rendahl KG, Karim C, Embry MG, Ghoddusi M, Holash J, et al. Neutralization of prolactin receptor function by monoclonal antibody LFA102, a novel potential therapeutic for the treatment of breast cancer. Molecular cancer therapeutics. 2013;12(3):295–305.

36. Kataoka S. SJ, Fujiya H., Toyota T., Suzuki R., Itoh K., Kumagai K. . Immunologic Aspects of the Nonobese Diabetic (NOD) Mouse: Abnormalities of Cellular Immunity Diabetes 1983;32(3):247–53.

37. Peiris-Pagès M, Federica, S., Lisanti, M.P. . Doxycycline and therepeutic targeting of DNA damage response in cancerl cells: old drug, new purpose Oncoscience 2015;2(8):696–9.

38. Kataoka S. KY, Nishio Y., Fujikawa-Adachi K., Tominaga A. Antitumor Activity of Eosinophils Activated by IL-5 and Eotaxin against Hepatocellular Carcinoma DNA and Cell Biology 2004;23(9):549–60.

39. Slifka MK, Pagarigan RR, Whitton JL. NK Markers Are Expressed on a High Percentage of Virus-Specific CD8+ and CD4+ T Cells. The Journal of Immunology. 2000;164(4):2009–15.

40. Trambley J, Bingaman AW, Lin A, Elwood ET, Waitze SY, Ha J, et al. Asialo GM1(+) CD8(+) T cells play a critical role in costimulation blockade-resistant allograft rejection. J Clin Invest. 1999;104(12):1715–22.

41. Nishikado H, Mukai K, Kawano Y, Minegishi Y, Karasuyama H. NK cell-depleting anti-asialo GM1 antibody exhibits a lethal off-target effect on basophils in vivo. J Immunol. 2011;186(10):5766–71.

42. Wiltrout RH SA, Peterson ES, Knott DC, Overton WR, et al. Reactivity of anti-asialo GM1 serum with tumouricidal and non-tumouricidal mouse macrophages. Journal of leukocyte biology 1985;37:597–614.

43. Dorshkind K, Pollack, S. B, Bosma, M.J., Philips, R.A. . Natural Killer (NK) cells are present in mice with severe combined immunodeficiency (scid) The Journal of Immunology. 1985;134:3798–801.

44. Yang Q. GSR, Hoklan M.E., Basse P.H. . Antitumor activity of NK cells. Immunologic Research 2006;36/1(3):13–25.

45. R.I T. The eosinophil-mediated antitumor activity of interleukin-4. J Allergy Clin Immunol 1994;94(1225-1230).

46. Sektioglu IM, Carretero R, Bulbuc N, Bald T, Tuting T, Rudensky AY, et al. Basophils Promote Tumor Rejection via Chemotaxis and Infiltration of CD8+ T Cells. Cancer research. 2017;77(2):291–302.

47. Chan CJ, Smyth MJ, Martinet L. Molecular mechanisms of natural killer cell activation in response to cellular stress. Cell death and differentiation. 2014;21(1):5–14.

48. Soriani A, Zingoni, A., Cerboni, C., Iannitto, M.L., Ricclardi, M.R., et al. ATM-ATR-dependent up-regulation of DNAM-1 and NKG2D ligands on multiple myloma cells by therapeutic agents results in enhanced NK-cell susceptibility and is associated with a senescent phenotype Blood. 2009;113(15):3503–11.

49. Wennerberg E, Sarhan D, Carlsten M, Kaminskyy VO, D’Arcy P, Zhivotovsky B, et al. Doxorubicin sensitizes human tumor cells to NK cell- and T-cell-mediated killing by augmented TRAIL receptor signaling. Int J Cancer. 2013;133(7):1643–52.

50. Dill R, Walker AM. Role of Prolactin in Promotion of Immune Cell Migration into the Mammary Gland. Journal of mammary gland biology and neoplasia. 2017;22(1):13–26.

51. Sun R. LAL, Wei H.M., Tian Z.G. . Expression of prolactin receptor and response to prolactin stimulation of human NK cells Cell Research 2004;14(1):67–73.

52. Sun R. WH, Zhang J., Li A., Zhang W., Tian Z.G. Recombinant human prolactin improves antitumor effecst of murine natural killer cells in vitro and in vivo Neuroimmunomodulation. 2002;10:169–76.

53. Zhang J, Sun R, Wei H, Tian Z. Antitumor effects of recombinant human prolactin in human adenocarcinoma-bearing SCID mice with human NK cell xenograft. Int Immunopharmacol. 2005;5(2):417–25.

54. Mavoungou E, Bouyou-Akotet MK, Kremsner PG. Effects of prolactin and cortisol on natural killer (NK) cell surface expression and function of human natural cytotoxicity receptors (NKp46, NKp44 and NKp30). Clin Exp Immunol. 2005;139(2):287–96.

