## Supplemental files for "Prolactin and DNA damage trigger an anti-breast cancer cell immune response"

### Supplementary Materials and Methods

#### Polymerase Chain Reactions

RNA was extracted from breast cancer cells using a Qiagen RNEasy Mini Kit (Qiagen Inc., Mississauga, ON, Canada) according to the manufacturer's protocol. Complimentary DNA (cDNA) was synthesized from 2 ug of RNA using a Superscript II Reverse Transcriptase kit (Invitrogen) according to the manufacturer's protocol.

All primers were designed using the NCBI Primer Blast-program. The Operon Oligo Analysis tool was used to detect possible primer dimers and self-complementation was identified with IDT Oligo Analyzer. The desired primers were obtained from the University of Calgary DNA Synthesis Lab (Calgary, AB). Quantitative polymerase chain reactions (qPCR) were carried out using iTaq Universal SYBR Green Supermix (Biorad, Mississauga, ON, Canada) with 1 $\mu$l of each forward and reverse primer (final primer concentration of 200 mM) (Table 1). The following protocol was performed for amplification of Sh ble gene: 95 °C for 2 minutes, 40 cycles of 95 °C for 5 seconds, 60 °C for 30 seconds, 78 °C for 20 seconds, and final extension step of 72 °C for 10 minutes, and following protocol was used for amplification of YWHAZ gene: 95 °C for 2 minutes, 40 cycles of 95 °C for 10 seconds, 60 °C for 30 seconds, 78 °C for 20 seconds, and final extension step of 72 °C for 10 minutes in the BioRad MJMini Opticon Real-Time PCR System.

#### Whole cell extract

$10^6$ cells were plated in 10cm plates, and the next day cells were washed with 1X PBS and directly scraped in 1X SDS protein sample buffer , followed by sonication three times for 5 seconds with 5-second intervals at #5 (on dial) (Fisher

Scientific 60 Sonic Dismembrator) on ice. Protein samples were snap frozen and stored at -80 °C until use.

**Supplementary Figures**

**S1. Confirmation of PRL-secreting and EV-control colonies A.** PRL levels were measured from whole cell extracts of MCF7 parental cells and MCF7hPRL colonies. GRB2 was used as a loading control. **B.** Western analysis of PRL secretion from MCF7hPRL colonies (1 to 5) compared with human recombinant PRL standard. **C.** PCR data showing Sh ble zeocin resistance gene from MCF7hPRL and MCF7EV colonies. *YWHAZ* was used as a house-keeping gene.

**S2. Autocrine PRL delays tumour latency in the presence of DNA damage in SCID mice**. **A**. Tumour latency in SCID mice after injection of 250,000 MCF7 or MCF7hPRL cells -/+ doxorubicin over 120 days. The sample size is n=5 mice for each group. Log-rank (Mantel-Cox) and Gehan-Breslow Wilcoxon tests were used for statistical analysis.

**S3. Cell viability (Alamar blue) assay showing anti-asialo GM1 does not affect the viability of MCF7, MCF7EV and MCF7hPRL cells in the presence and absence of doxorubicin.** Cells were treated with 20 ul/2.5 ml, 20 ul/2 ml, 20 ul/1.5ml, 20 ul/1.25 ml of anti-asialo GM1. The cell viability was followed 96 hrs. Graphs represent pooled experiments, n=6. **A.** Cell viability in MCF7, MCF7EV and MCF7hPRL cells with anti-asialo GM1 treatment after 24hrs. **B.** Cell viability in MCF7, MCF7EV and MCF7hPRL cells with anti-asialo GM1 treatment after 48 hrs **C.** Cell viability in MCF7, MCF7EV and MCF7hPRL cells with anti-asialo GM1 treatment after 73 hrs **D.** Cell viability in MCF7, MCF7EV and MCF7hPRL cells with/without human recombinant prolactin (25 ng/ml), doxorubicin (1 uM) and anti-asialo GM1 treatment after 24 hrs. **E.** Cell viability in MCF7, MCF7EV and MCF7hPRL cells with/ without human recombinant prolactin (25 ng/ml), doxorubicin (1 uM) and anti-asialo GM1 treatment after 48 hrs. **F.** Cell viability in MCF7,

MCF7EV and MCF7hPRL cells with/without human recombinant prolactin (25 ng/ml), doxorubicin (1 uM) and anti-asialo GM1 treatment after 72 hrs.

**S4. Calcein-AM assay determining the NK cell-mediated lysis of MCF7, MCF7hPRL and MC7EV cells in the presence or absence of DNA damage.** MCF7 and MCF7EV cells were pre-treated with human recombinant prolactin (25 ng/ml) for 24 hours, followed by 2 hours of doxorubicin treatment (1 uM). Cells were trypsinized after 48 hours of recovery time and co-cultured with NK cells in a 1:1 or 10:1 effector/ target ratio. The cell viability of breast cancer cells was determined by Calcein-AM assay and the % Lysis was calculated. **A**. MCF7 (1:1 E/F ratio). **B**. MCF7EV (1:1 E/F ratio). **C**. MCFhPRL (1:1 E/F ratio). D. MCF7 (10:1 E/F ratio). E. SKBR3 (10:1 E/F ratio) Statistically significant analysis (*) denotes *P*<.05, (**) denotes *P*<.01, (***) denotes *P*<.001.

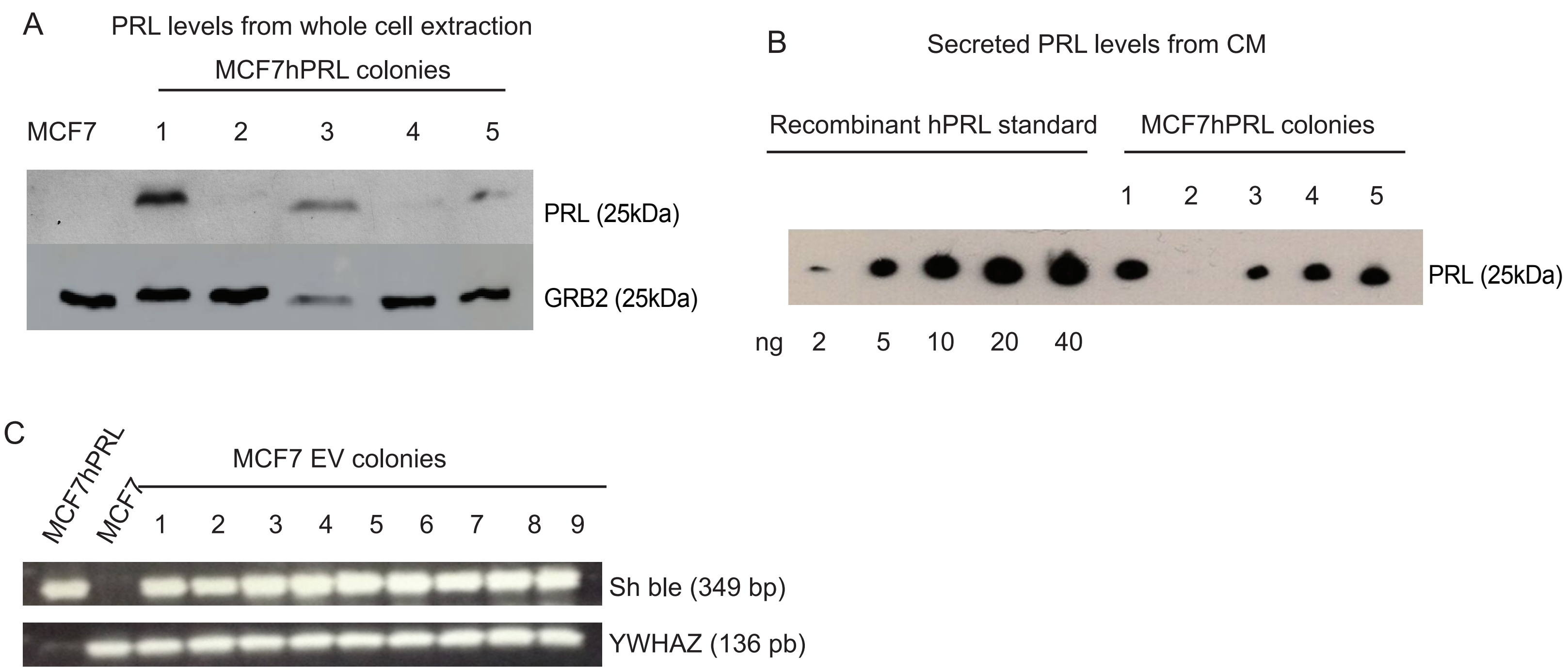
A
PRL levels from whole cell extraction
MCF7hPRL colonies
MCF7
1
2
3
4
5
PRL (25kDa)
GRB2 (25kDa)
B
Secreted PRL levels from CM
Recombinant hPRL standard
MCF7hPRL colonies
1
2
3
4
5
PRL (25kDa)
ng
2
5
10
20
40
C
MCF7hPRL
MCF7
MCF7 EV colonies
1
2
3
4
5
6
7
8
9
Sh ble (349 bp)
YWHAZ (136 pb)

Figure S1

A

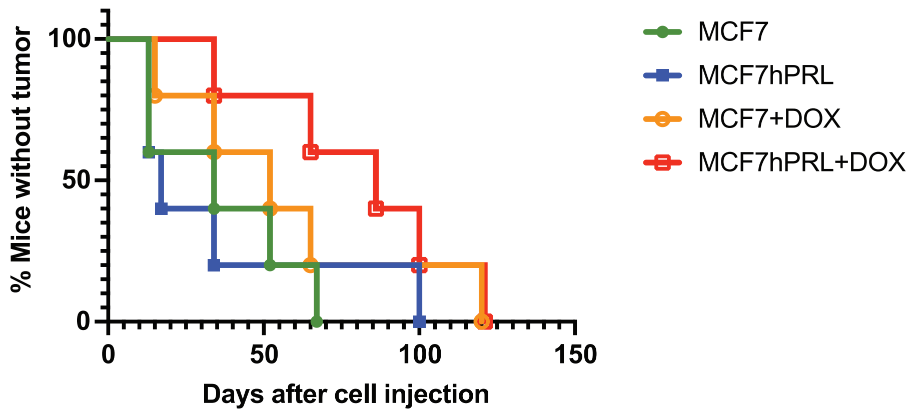

% Mice without tumor
100
50
0
0
50
100
150
Days after cell injection
MCF7
MCF7hPRL
MCF7+DOX
MCF7hPRL+DOX


Figure S2

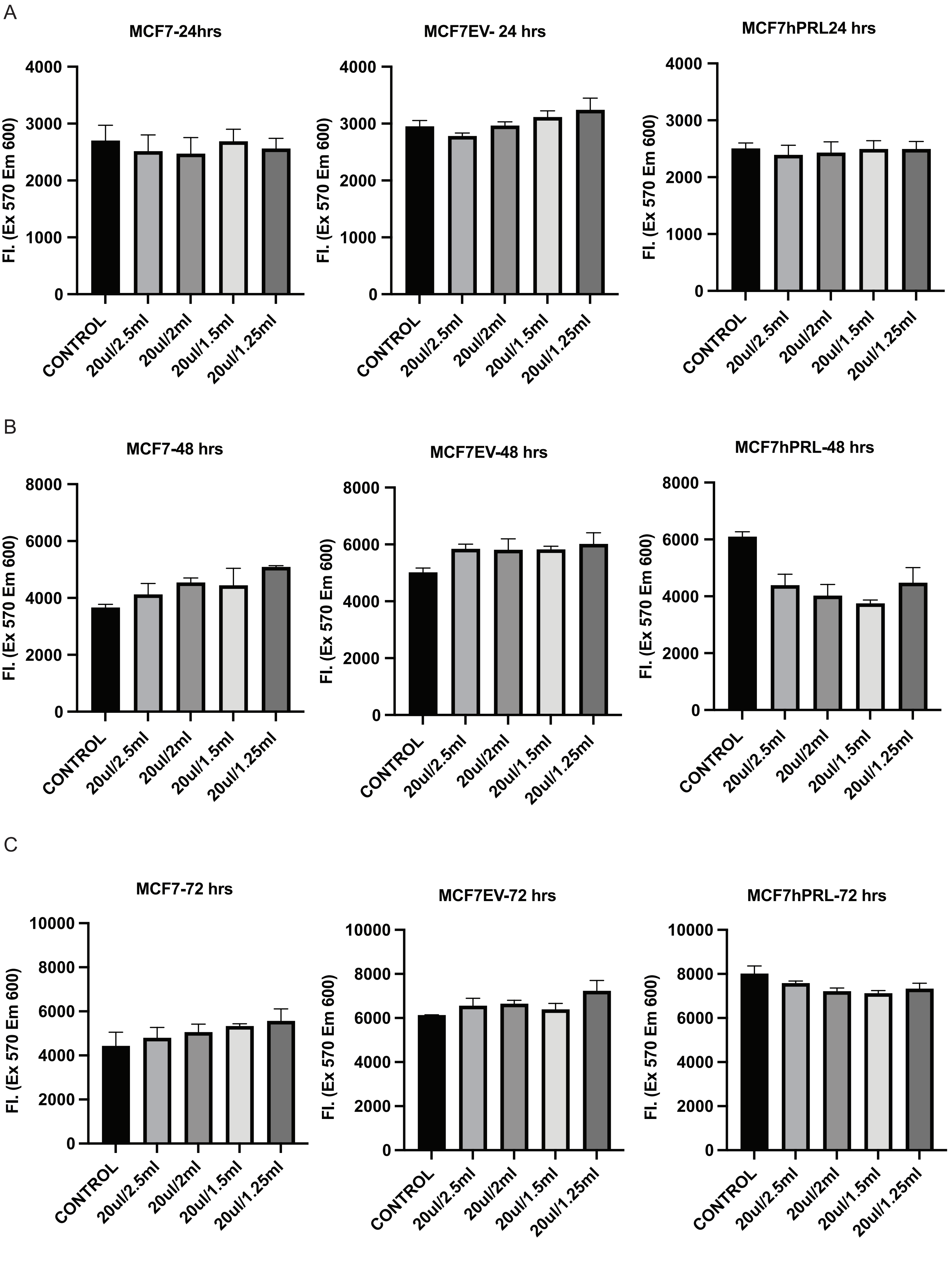
A
MCF7-24hrs
MCF7EV- 24 hrs
MCF7hPRL24 hrs
Fl. (Ex 570 Em 600)
0
1000
2000
3000
4000
CONTROL
20ul/2.5ml
20ul/2ml
20ul/1.5ml
20ul/1.25ml
B
MCF7-48 hrs
MCF7EV-48 hrs
MCF7hPRL-48 hrs
Fl. (Ex 570 Em 600)
0
2000
4000
6000
8000
CONTROL
20ul/2.5ml
20ul/2ml
20ul/1.5ml
20ul/1.25ml
C
MCF7-72 hrs
MCF7EV-72 hrs
MCF7hPRL-72 hrs
Fl. (Ex 570 Em 600)
0
2000
4000
6000
8000
10000
CONTROL
20ul/2.5ml
20ul/2ml
20ul/1.5ml
20ul/1.25ml

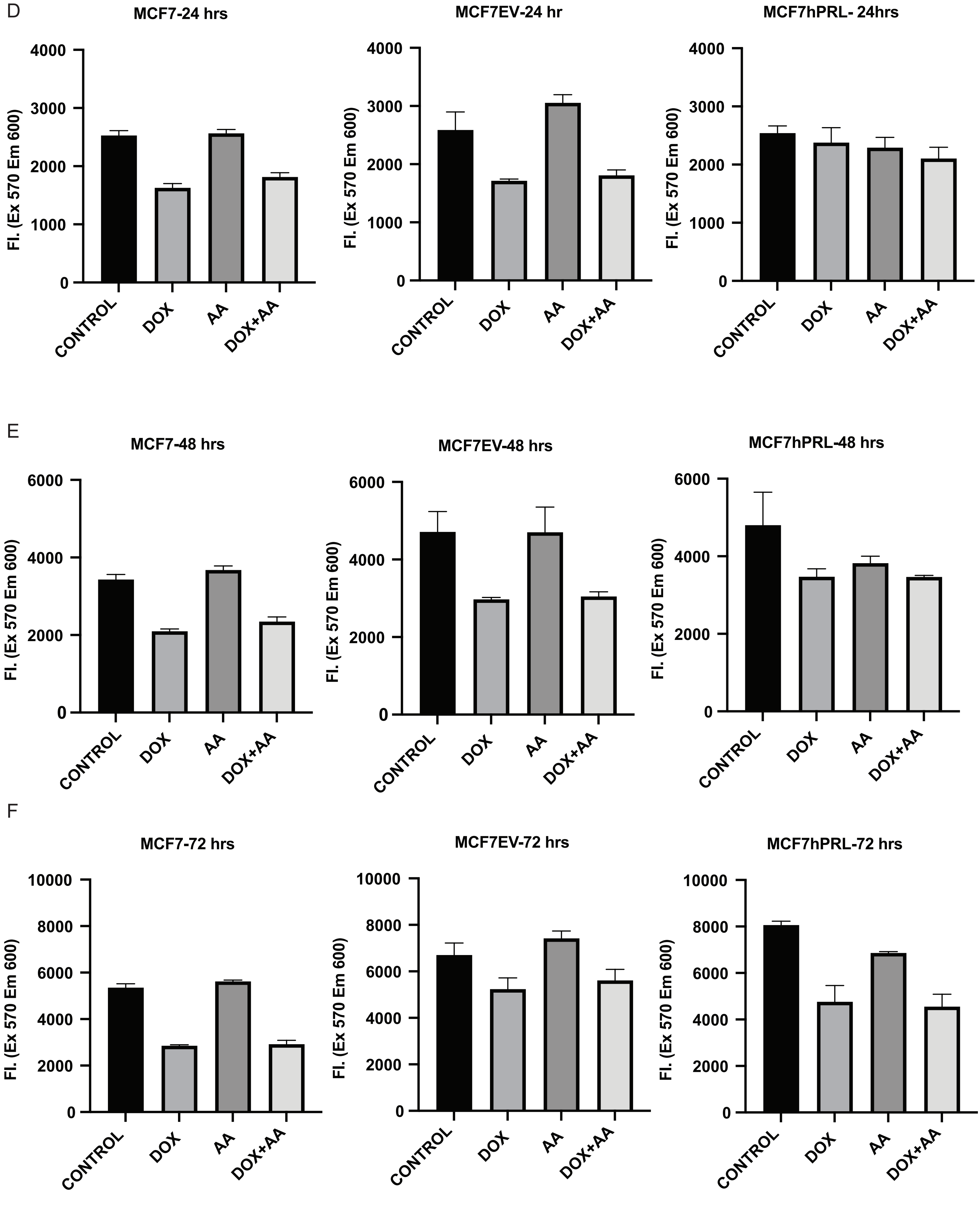
D
MCF7-24 hrs
MCF7EV-24 hr
MCF7hPRL- 24hrs
Fl. (Ex 570 Em 600)
0
1000
2000
3000
4000
CONTROL
DOX
AA
DOX+AA
E
MCF7-48 hrs
MCF7EV-48 hrs
MCF7hPRL-48 hrs
0
2000
4000
6000
F
MCF7-72 hrs
MCF7EV-72 hrs
MCF7hPRL-72 hrs
0
2000
4000
6000
8000
10000

Figure S3

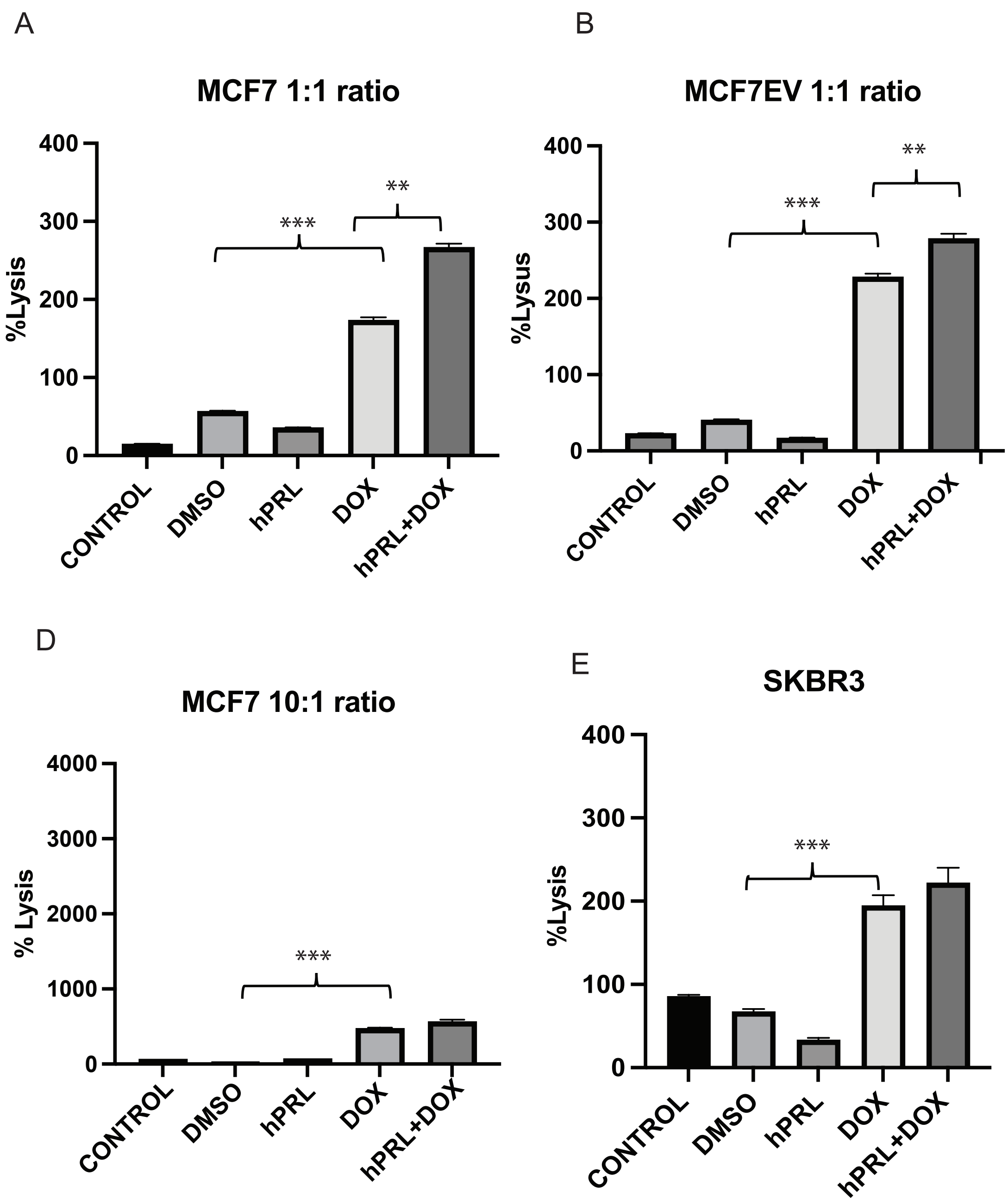
A
MCF7 1:1 ratio
%Lysis
400
300
200
100
0
***
**
CONTROL
DMSO
hPRL
DOX
hPRL+DOX
B
MCF7EV 1:1 ratio
%Lysus
400
300
200
100
0
***
**
CONTROL
DMSO
hPRL
DOX
hPRL+DOX
C
MCF7hPRL 1:1 ratio
%Lysus
400
300
200
100
0
***
CONTROL
DMSO
DOX
D
MCF7 10:1 ratio
% Lysis
4000
3000
2000
1000
0
***
CONTROL
DMSO
hPRL
DOX
hPRL+DOX
E
SKBR3
%Lysis
400
300
200
100
0
***
CONTROL
DMSO
hPRL
DOX
hPRL+DOX

Figure S4

**Supplementary Table 1 Primers**

| **Primer name** | **Primers** | **Tm (ºC)** | **Expected amplicon size (pb)** |
|---|---|---|---|
| Steptoalloteichus hindustanus bleomycin (Sh-ble) (Zeocin resistance gene) | F- 5”-AAGTTGACCAGTGCCGTTCC-3’<br>R- 5’-CTCCTCGGCCACGAAGTG-3’ | 60° C | 360 |
| Tyrosine 3-monooxygenase/tryptophan 5-monooxygenase activation protein (YWHAZ) | F- 5’-AGTCGTACAAAGACAGCACGTAA-3’<br>R- 5’-AGGCAGACAAAGGTTGGAAGG-3’ | 60° C | 138 |
